# Substrate-mediated regulation of the arginine transporter of *Toxoplasma gondii*

**DOI:** 10.1101/798967

**Authors:** Esther Rajendran, Morgan Clark, Cibelly Goulart, Birte Steinhöfel, Erick T. Tjhin, Nicholas C. Smith, Kiaran Kirk, Giel G. van Dooren

## Abstract

Intracellular parasites, such as the apicomplexan *Toxoplasma gondii*, are adept at scavenging nutrients from their host. However, there is little understanding of how parasites sense and respond to the changing nutrient environments they encounter during an infection. *Tg*ApiAT1, a member of the apicomplexan ApiAT family of amino acid transporters, is the major uptake route for the essential amino acid L-arginine (Arg) in *T. gondii*. Here, we show that the abundance of *Tg*ApiAT1, and hence the rate of uptake of Arg, is regulated by the availability of Arg in the parasite’s external environment, increasing in response to decreased [Arg]. Using a luciferase-based ‘biosensor’ strain of *T. gondii*, we demonstrate that parasites vary the expression of *Tg*ApiAT1 in different organs within their host, indicating that parasites are able to modulate *Tg*ApiAT1-dependent uptake of Arg as they encounter different nutrient environments *in vivo*. Finally, we show that Arg-dependent regulation of *Tg*ApiAT1 expression is post-transcriptional, mediated by an upstream open reading frame (uORF) in the *Tg*ApiAT1 transcript, and we provide evidence that the peptide encoded by this uORF is critical for mediating regulation. Together, our data reveal the mechanism by which an apicomplexan parasite responds to changes in the availability of a key nutrient.

## INTRODUCTION

Apicomplexans are a phylum of intracellular parasites that include the causative agents of malaria (*Plasmodium* spp.) and toxoplasmosis (*Toxoplasma gondii*). The proliferation of parasites in their hosts, and their progression through their often complex life cycles, is dependent on nutrients scavenged from the host [1–3]. Apicomplexans encounter different nutrient conditions as they proliferate within, and move between, hosts, and this is reflected in differences in the metabolism of different parasite life-stages; *e.g.*, hepatocyte stages of *Plasmodium* parasites rely on the biosynthesis of haem and fatty acids, whereas the intra-erythrocytic parasite stages scavenge these from the host [4, 5]. Although early studies suggested that parasite metabolism is ‘hard-wired’ and resistant to adapting to changes in nutrient conditions [6], there is growing evidence that parasites sense and respond to changes in the nutrient status of their hosts [3]. For example, *Plasmodium* blood-stage parasites modulate their proliferation in response to the caloric intake of their hosts, and can enter a dormant state in response to limitation of the essential amino acid isoleucine [7, 8].

In some instances, the ability of parasites to sense changes in external nutrient levels is key to their differentiation into new life stages. For example, limitation of lysophosphatidylcholine induces *Plasmodium falciparum* parasites to differentiate into the transmitted sexual stages in the human host [9], and the high levels of linoleic acid that *T. gondii* parasites encounter in the intestines of felids induces parasite differentiation into the sexual stages [10]. The depletion of the amino acid arginine (Arg), which may be caused by host immune responses [11], is thought to lead to differentiation of the disease-causing tachyzoite stage of *T. gondii* into the dormant, cyst-forming bradyzoite stage [12]. Despite the importance of nutrient sensing in parasite proliferation and differentiation, the mechanisms by which parasites sense and respond to the availability of nutrients are largely unknown.

The uptake of nutrients by *T. gondii* parasites is mediated primarily by plasma membrane transporters [13]. We recently characterised a family of plasma membrane amino acid transporters that are found throughout apicomplexans and have termed these the Apicomplexan Amino acid Transporter (ApiAT) family [14]. We have demonstrated that one member of this family, *Tg*ApiAT1, is an Arg transporter that is essential for *T. gondii* virulence [15].

In this study, we have investigated the ability of *T. gondii* parasites to sense and respond to the Arg levels that they encounter in their host. We report Arg-dependent regulation of *Tg*ApiAT1 expression, and demonstrate that this process is mediated by an upstream open reading frame (uORF) in the *Tg*ApiAT1 transcript. We also present evidence, obtained using a luciferase-based ‘biosensor’ strain of *T. gondii*, that parasites vary the expression of *Tg*ApiAT1 in different organs within their host. Our data demonstrate how *T. gondii* parasites are able to sense and respond to changes in the abundance of a key nutrient, as well as illustrating their ability to do so within the course of an infection.

## RESULTS

### Regulation of *Tg*ApiAT1 protein abundance and parasite arginine uptake

To investigate whether the abundance of *Tg*ApiAT1 is dependent upon Arg availability, we introduced a haemagglutinin (HA_3_) epitope tag into the open reading frame of the *Tg*ApiAT1 genomic locus. The resultant *Tg*ApiAT1-HA_3_-expressing parasites were grown in modified Roswell Park Memorial Institute 1640 (RPMI) medium in which [Arg] ranged from 10 µM to 5 mM. Western blotting revealed that the abundance of *Tg*ApiAT1-HA_3_ varied with [Arg], with *Tg*ApiAT1-HA_3_ most abundant in parasites grown at low [Arg] (Figure 1A).

**Figure 1.**
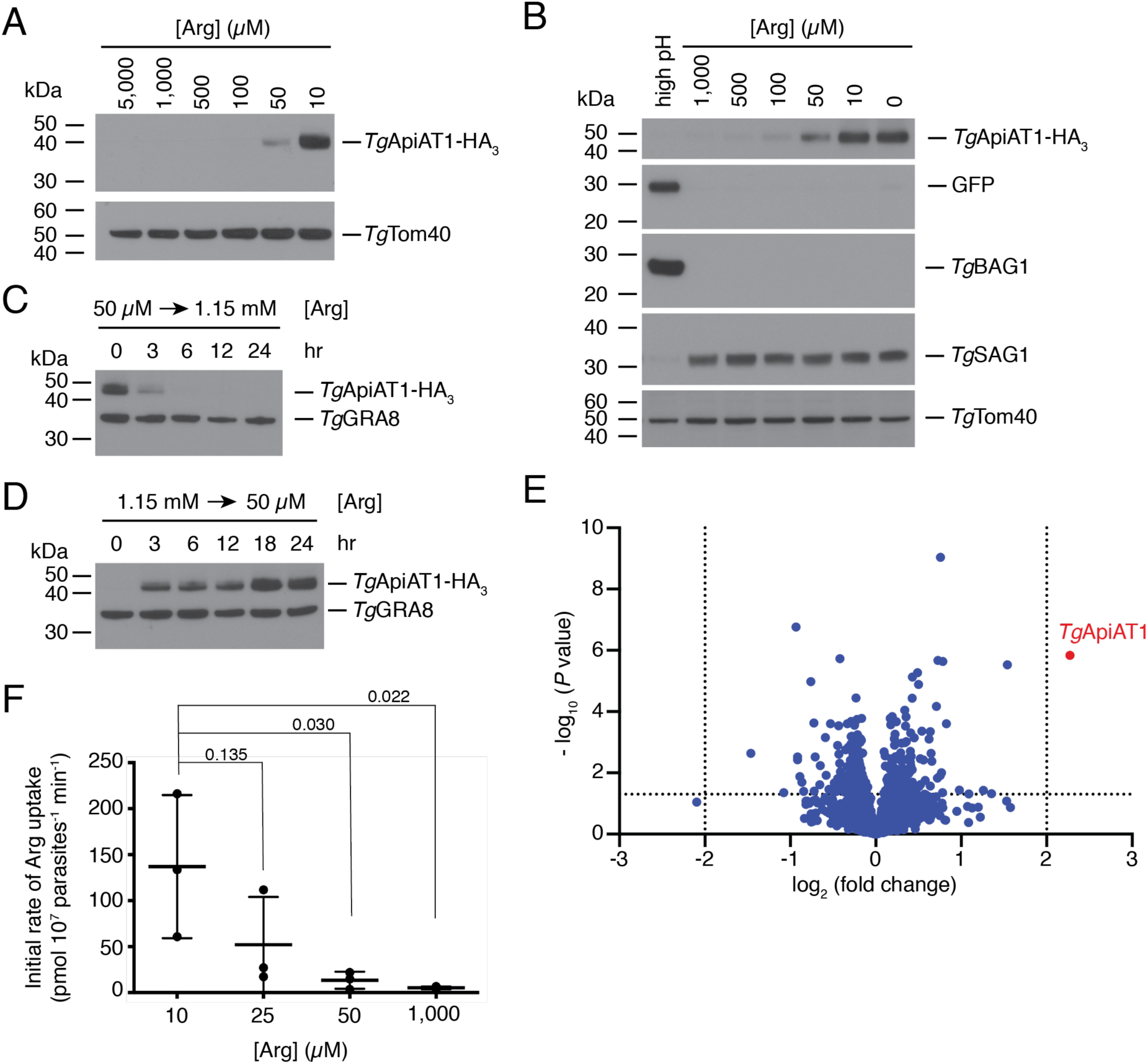
*Tg*ApiAT1 protein abundance is regulated by [Arg] in the growth medium. (**A**) Western blot of *Tg*ApiAT1-HA_3_ in parasites grown at a range of [Arg] in the growth medium. *Tg*Tom40 is a loading control. Data are representative of three independent experiments. (**B**) Western blot of *Tg*ApiAT1-HA_3_ in Prugniaud strain parasites co-expressing GFP from the bradyzoite-specific *Tg*LDH2 upstream region. Parasites were grown at a range of [Arg] in the growth medium, or a high pH to induce bradyzoite formation, and probed with antibodies against anti-HA (to detect *Tg*ApiAT1-HA_3_), anti-GFP (to detect GFP expressed from the bradyzoite-specific promoter LDH2), anti-BAG1 (a bradyzoite-specific marker), anti-SAG1 (a tachyzoite-specific marker), and anti-*Tg*Tom40 (a loading control). (**C, D**) Western blot of *Tg*ApiAT1-HA_3_ in parasites grown at low (50 µM; **C**) or high (1.15 mM; **D**) [Arg] and switched to high or low [Arg], respectively, for the indicated times. *Tg*GRA8 is a loading control. Data are representative of three independent experiments. (**E**) Volcano plot depicting log_2_ fold change vs -log_10_ *P* values of change in protein abundance in a SWATH MS-based proteomic analysis of parasites grown at 50 µM vs 1.15 mM Arg (n = 5). Dotted lines represent values where *P* = 0.05 (y axis) or log_2_ fold change is −2 or 2 (x axis). The *Tg*ApiAT1 data point is depicted in red. (**F**) Initial rate of Arg uptake in parasites cultured in growth medium containing 10, 25, 50 or 1,000 µM Arg. Uptake was measured in 50 µM unlabelled Arg and 0.1 µCi/ml [^14^C]Arg. Initial rates were calculated from the initial slope of fitted single-order exponential curves of timecourse uptake experiments (Figure S1). Data represent the mean ± SD from three independent experiments. *P* values were calculated using a one-way ANOVA with Dunnett’s multiple comparisons test, comparing the initial uptake rates to the 10 µM condition.

Low [Arg] conditions have been linked to formation of the latent bradyzoite stage of *T. gondii* [12]. We measured *Tg*ApiAT1-HA_3_ abundance in Type II Prugniaud strain *T. gondii* parasites, which readily form bradyzoites, and observed regulation of *Tg*ApiAT1-HA_3_ levels in response to variation in [Arg] but no variation of *Tg*ApiAT1-HA_3_ levels in response to pH-mediated bradyzoite induction (Figure 1B). This indicates that *Tg*ApiAT1 regulation is not related to the parasite’s general bradyzoite differentiation response.

To assess the kinetics of the Arg-dependence of *Tg*ApiAT1-HA_3_ expression, we switched *Tg*ApiAT1-HA_3_ parasites grown at 50 µM Arg to medium containing 1.15 mM Arg for 3-24 hr. *Tg*ApiAT1-HA_3_ protein levels decreased within 3 hr of the switch (Figure 1C). In the converse experiment, when *Tg*ApiAT1-HA_3_ parasites were switched from medium containing 1.15 mM Arg to medium containing 50 µM Arg, *Tg*ApiAT1-HA_3_ protein levels increased within 3 hr of the switch (Figure 1D). These data reveal that *T. gondii* parasites change the abundance of their major Arg transporter in response to the [Arg] they encounter in their growth medium, doing so within hours.

To assess whether the abundance of other proteins changed upon changes to [Arg] in the growth medium, we grew parasites in media containing either 50 µM or 1.15 mM Arg and extracted proteins for quantitative proteomics using sequential window acquisition of all theoretical fragment ion spectra mass spectrometry (SWATH-MS; [16]). This revealed that *Tg*ApiAT1 was the only protein for which abundance was significantly increased beyond a log_2_ fold change of 2 in the 50 µM compared to the 1.15 mM condition (Figure 1E; Table S1; *P* < 0.05).

To establish whether changes in *Tg*ApiAT1 abundance correlate with changes in Arg uptake by the parasite, we grew parasites at a range of [Arg] and measured *Tg*ApiAT1-dependent [^14^C]-labelled Arg uptake. In parasites grown at 10 µM Arg the initial rate of Arg uptake was 25-fold higher than in parasites grown at 1 mM Arg (Figure 1F; Figure S1).

### The 5’ region of the *Tg*ApiAT1 gene regulates *Tg*ApiAT1 protein abundance

The expression of many proteins is mediated by genetic information encoded upstream (5’) of the start codon. To test whether the 5’ region of the gene encoding *Tg*ApiAT1 is important for regulation, we measured *Tg*ApiAT1-HA_3_ abundance in a strain in which *Tg*ApiAT1-HA_3_ was expressed from the *α*-tubulin promoter and in which the native *Tg*ApiAT1 gene had been knocked out [15]. We grew this strain at 10 µM, 50 µM and 1 mM Arg. Western blotting revealed no variation in *Tg*ApiAT1-HA_3_ abundance (Figure 2A), indicating that the 5’ region of the *Tg*ApiAT1 coding sequence is necessary for Arg-dependent regulation of *Tg*ApiAT1.

**Figure 2.**
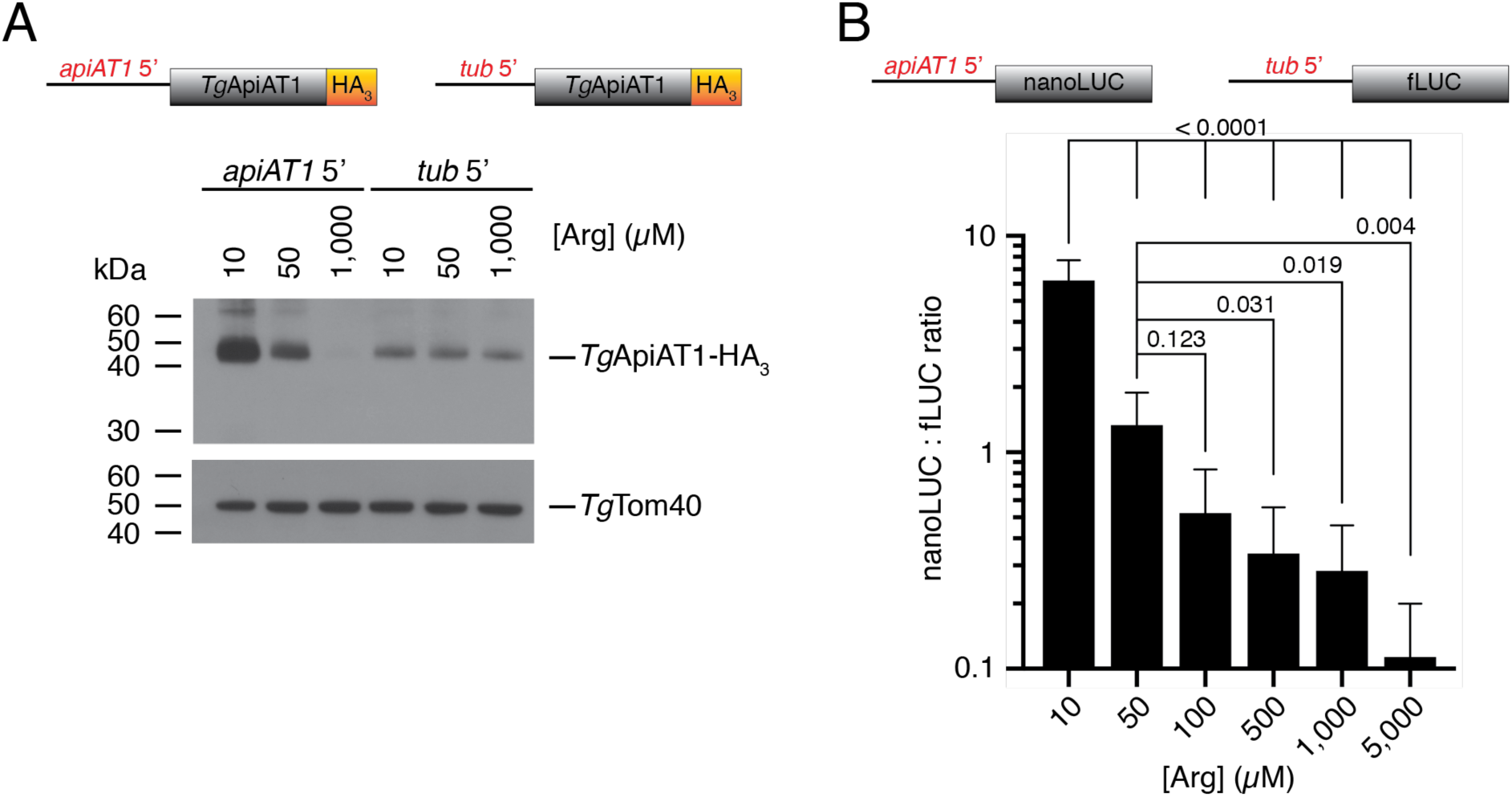
Arg-dependent *Tg*ApiAT1 regulation is mediated by the 5’ upstream region of the *Tg*ApiAT1 gene. (**A**) Western blot of *Tg*ApiAT1-HA_3_ expressed from the native *Tg*ApiAT1 5’ region (*apiAT1* 5’) or the *α*-tubulin 5’ region (*tub* 5’), in parasites grown at a range of [Arg] in the growth medium. *Tg*Tom40 is a loading control. Data are representative of three independent experiments. (**B**) nanoLUC:fLUC ratio in a parasite strain expressing nanoLUC from the *Tg*ApiAT1 5’ region (*apiAT1* 5’-nanoLUC) and fLUC from the *α*-tubulin 5’ region (*tub* 5’-fLUC), and grown at a range of [Arg]. Data represent the mean ± SD from nine independent experiments. *P* values were calculated using a one-way ANOVA with Tukey’s multiple comparisons test. *P* values not shown were > 0.500.

To determine whether the 5’ region of the gene encoding *Tg*ApiAT1 is sufficient to mediate Arg-dependent regulation, we expressed a nanoLUC luciferase (nanoLUC) reporter enzyme from the *Tg*ApiAT1 5’ region in a strain that expressed a firefly luciferase (fLUC) reporter from the *α*-tubulin 5’ region (Figure 2B). We grew this ‘dual reporter’ strain at [Arg] ranging from 10 µM to 5 mM and measured nanoLUC- and fLUC-dependent luminescence. NanoLUC-dependent luminescence decreased with increasing [Arg], whereas fLUC-dependent luminescence remained unchanged (Figure S2). This enabled fLUC luminescence to be used as a normalising factor, with the nanoLUC:fLUC luminescence ratio providing a measure of Arg-dependent regulation mediated by the 5’ region of the gene encoding *Tg*ApiAT1. There was a significant decrease in the nanoLUC:fLUC ratio as [Arg] increased, with a 55-fold decrease in parasites grown at 5 mM Arg relative to parasites grown at 10 µM Arg (Figure 2B). Expression of nanoLUC from the *α*-tubulin 5’ region revealed no Arg-dependent regulation (Figure S2), ruling out the possibility that nanoLUC expression is itself Arg-dependent. We conclude that the 5’ region of the gene encoding *Tg*ApiAT1 is both *necessary* and *sufficient* to mediate Arg-dependent regulation of the *Tg*ApiAT1 protein.

### *Tg*ApiAT1 abundance is regulated by the availability of other nutrients, including lysine, in an opposite manner to arginine

Next, we asked whether *Tg*ApiAT1 expression is regulated by the availability of other nutrients. We measured the abundance of *Tg*ApiAT1-HA_3_ in parasites grown in media containing from 62.5 µM to 1 mM L-lysine (Lys) at a constant 50 µM Arg. We observed increased protein abundance with increased [Lys] (Figure 3A), the *opposite* effect to what was observed with changing [Arg]. Similarly, when the dual reporter strain was grown in media ranging from 62.5 µM to 1 mM Lys and a constant 50 µM Arg, the nanoLUC:fLUC ratio increased with increasing [Lys] (Figure 3B; Figure S3A). To investigate this further, we measured the nanoLUC:fLUC luminescence ratio at a range of [Arg] at high (1 mM) or low (50 µM) Lys. At all but the lowest Arg concentration tested (*i.e.* 10 µM), the nanoLUC:fLUC luminescence ratio measured in parasites grown at high [Lys] was higher than that measured in parasites grown at low [Lys] (Figure 3C). Together these results indicate that [Lys] influences *Tg*ApiAT1 expression in the *opposite* manner to [Arg].

**Figure 3.**
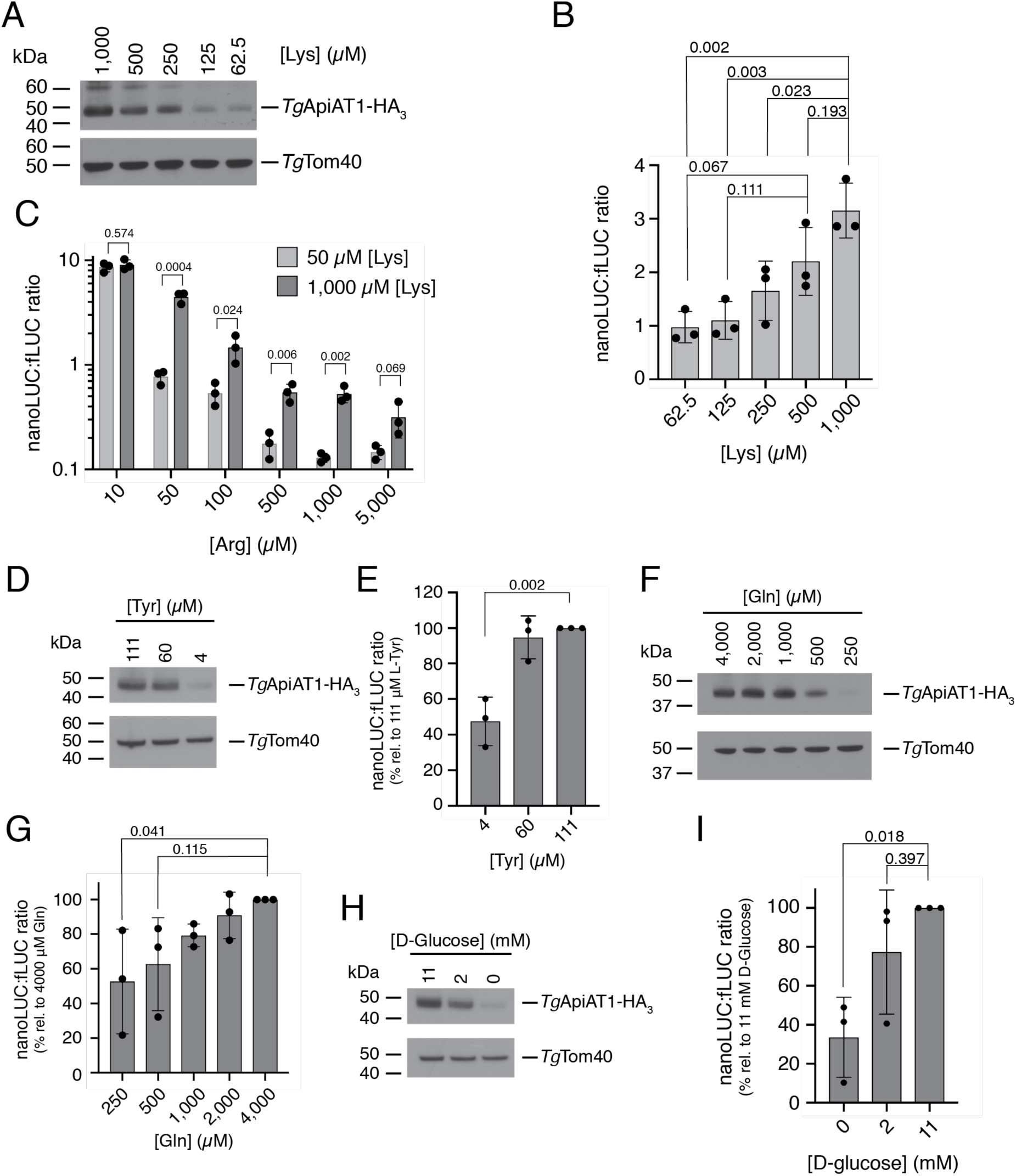
*Tg*ApiAT1 regulation is mediated by a range of nutrients. (**A**, **D**, **F**, **H**) Western blots of *Tg*ApiAT1-HA_3_ in parasites grown at a range of (**A**) [Lys], (**D**) [Tyr], (**F**) [Gln], or (**H**) [D-glucose] in the growth medium. *Tg*Tom40 is a loading control. Data are representative of three independent experiments. (**B**) nanoLUC:fLUC ratio in parasites grown in media containing a range of concentrations of Lys. Data represent the mean ± SD from three independent experiments. *P* values were calculated using a one-way ANOVA with Tukey’s multiple comparisons test. *P* values not shown were > 0.400. (**C**) nanoLUC:fLUC ratios in parasites grown in media containing a range of [Arg] and either 50 µM Lys (light grey) or 1 mM Lys (dark grey). Data represent the mean ± SD from three independent experiments. *P* values were calculated using unpaired t-tests, not assuming equal variance (d.f. = 4). (**E**, **G**, **I)** nanoLUC:fLUC ratios in parasites grown in media containing a range of concentrations of (**E**) Tyr, (**G**) Gln, or (**I**) D-glucose. Data represent the mean ± SD from three independent experiments, with the ratios normalised to the condition with the highest nutrient concentration. *P* values were calculated using a one-way ANOVA with Dunnett’s multiple comparisons test, comparing the normalised nanoLUC:fLUC ratios at each nutrient concentration to the condition containing the highest concentration tested. *P* values not shown were > 0.500.

We examined the effects of the concentration of a range of other nutrients, including L-tyrosine (Tyr), L-glutamine (Gln) and D-glucose, on the nanoLUC:fLUC luminescence ratio in the dual reporter strain and on *Tg*ApiAT1-HA_3_ protein abundance. At the lowest concentration of each nutrient tested, we observed decreased *Tg*ApiAT1-HA_3_ protein abundance and a significantly decreased nanoLUC:fLUC ratio (Figure 3D-I; *P* < 0.05; Figure S3B-D). The lowest concentrations tested for Tyr and Gln were close to the minimal amount of those nutrients required for optimal parasite growth [14, 17]. This is consistent with the hypothesis that *Tg*ApiAT1 abundance can be negatively regulated through a general amino acid starvation response in the parasite [17], and that this regulation is mediated by the 5’ upstream region of *Tg*ApiAT1. These hypotheses were not further investigated here.

The effect of [Lys] on the expression of *Tg*ApiAT1 was explored further. Our previous study revealed a connection between the uptake of Arg and Lys in *T. gondii*, demonstrating the presence of a cationic amino acid transporter that has a higher affinity for Lys than for Arg [15]. This transporter can take up sufficient Arg for parasite growth in the absence of *Tg*ApiAT1 if the concentration of Lys, a competitive inhibitor of Arg uptake via the transporter, is low [15]. Our unpublished research indicates that this transporter is *Tg*ApiAT6-1 (Figure 4A; Rajendran, Fairweather, *et al*., in preparation), another member of the *Tg*ApiAT family that localises to the parasite plasma membrane [14]. Using an HA-tagged *Tg*ApiAT6-1 strain [14], we asked whether *Tg*ApiAT6-1-HA_3_ abundance is regulated during growth in media containing a range of amino acid concentrations. We found that the abundance of *Tg*ApiAT6-1-HA_3_ did not differ in any of the tested Arg or Lys concentrations (Figure 4B-C) although, as for *Tg*ApiAT1-HA_3_, we did observe a decrease in protein abundance at low [Gln] (Figure 4D).

**Figure 4.**
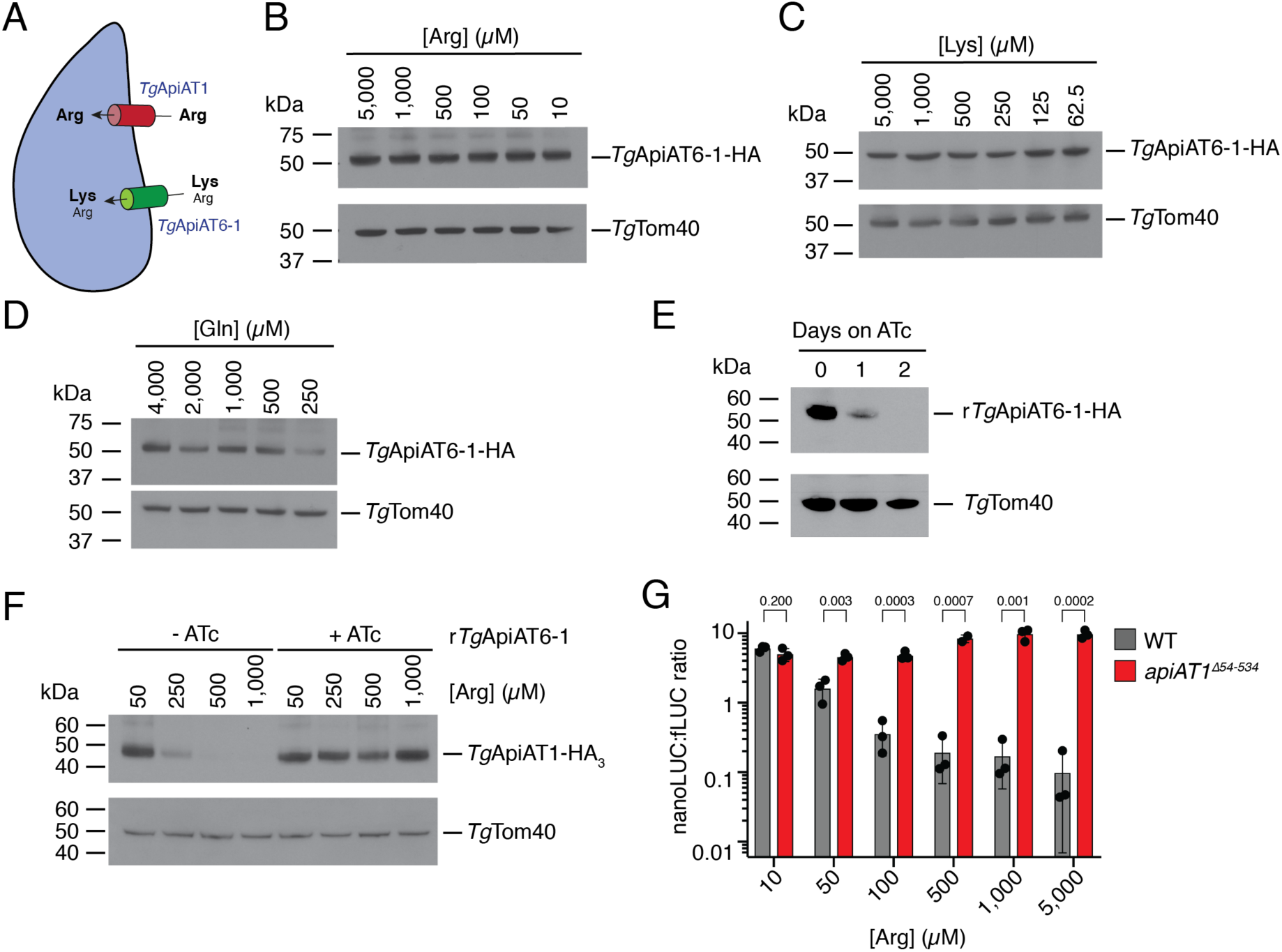
*Tg*ApiAT1 regulation is dependent on transporter-mediated uptake of Arg and Lys into the parasite. (**A**) Model of Arg uptake by *T. gondii*. *Tg*ApiAT1 is a selective Arg transporter, while *Tg*ApiAT6-1 is a cationic amino acid transporter with a high affinity for Lys and a lower affinity for Arg. (**B-D**) Western blots measuring the abundance of *Tg*ApiAT6-1-HA-expressing parasites grown at a range of (**B**) [Arg], (**C**) [Lys], or (**D**) [Gln] in the growth medium. *Tg*Tom40 is a loading control. Data are representative of two independent experiments. (**E**) Western blot measuring the abundance of r*Tg*ApiAT6-1-HA_3_ upon the addition ATc for 0 to 2 days. *Tg*Tom40 is a loading control. Data are representative of three independent experiments. (**F**) Western blot of *Tg*ApiAT1-HA_3_ in r*Tg*ApiAT6-1 parasites grown in the absence or presence of ATc, and at a range of [Arg] in the growth medium. *Tg*Tom40 is a loading control. Western blots are representative of three independent experiments. (**G**) nanoLUC:fLUC ratios in WT and *apiAT1*^Δ*54-534*^ parasites grown at a range of [Arg]. Data represent the mean ± SD from three independent experiments. *P* values were calculated using unpaired t-tests, not assuming equal variance (d.f. = 4). Note that the data from the WT experiments were also included in replicates for the data shown in Figure 2B.

The data from Figures 1 and 3 indicate that [Arg] and [Lys] have opposite effects on *Tg*ApiAT1 regulation. We considered two hypotheses to explain these data:

1. That *Tg*ApiAT1 regulation responds directly to [Lys] in the parasites.
2. That increased [Lys] in the growth medium results in increasing competition by lysine with arginine for the *Tg*ApiAT6-1 transporter. In turn, this leads to decreased uptake of [Arg] through *Tg*ApiAT6-1 and to lower [Arg] in the parasite, which subsequently results in an increase in *Tg*ApiAT1 expression. In this scenario, *Tg*ApiAT1 regulation responds only to [Arg] in the parasites.

To distinguish between the two possibilities, we generated a regulatable *Tg*ApiAT6-1 (r*Tg*ApiAT6-1) parasite strain, in which *Tg*ApiAT6-1 expression can be knocked down through the addition of anhydrotetracycline (ATc; Figure S4). We introduced a HA tag into the r*Tg*ApiAT6-1 strain and found that *Tg*ApiAT6-1-HA protein was undetectable after two days growth in ATc (Figure 4E). We then introduced a HA tag into the *Tg*ApiAT1 locus of the original r*Tg*ApiAT6-1 strain and grew parasites in the absence or presence of ATc at [Arg] ranging from 50 µM to 1 mM. *Tg*ApiAT1-HA_3_ abundance decreased with increasing [Arg] in the absence of ATc (in which *Tg*ApiAT6-1 is expressed) but remained invariant with varying [Arg] when ATc was added (and *Tg*ApiAT6-1 was depleted; Figure 4F). These data are consistent with the second hypothesis – that limiting Arg-uptake through *Tg*ApiAT6-1 leads to an increase in *Tg*ApiAT1 expression, and that Lys-dependent upregulation of *Tg*ApiAT1 (Figure 3A-C) results from reduced [Arg] in the parasite rather than increased [Lys].

In our previous study we found that knockout of *Tg*ApiAT1 led to decreased Arg uptake, which is expected to lead to reduced [Arg] in the parasite [15]. To explore further the relationship between parasite [Arg] and *Tg*ApAT1 regulation, we introduced a ‘knockout’ frameshift mutation in the *Tg*ApiAT1 locus of the dual luciferase reporter strain, generating a strain we termed *apiAT1*^Δ*54-534*^. As demonstrated previously for parasites lacking *Tg*ApiAT1 [14, 15], *apiAT1*^Δ*54-534*^ parasites exhibited reduced proliferation over an 8-day growth assay in Dulbecco’s modified Eagle’s medium (DME, which contains 400 µM Arg and 800 µM Lys) but grew normally in RPMI (which contains 1.15 mM Arg and 200 µM Lys) (Figure S5). We grew the *apiAT1*^Δ*54-534*^ strain in modified RPMI containing 10 µM to 5 mM Arg for 42 hr and measured the nanoLUC:fLUC luminescence ratio. In contrast to WT parasites, the nanoLUC:fLUC ratio in the *apiAT1*^Δ*54-534*^ strain did not decrease with increasing [Arg] (Figure 4G).

Taken together, the data from Figure 4 indicate that Arg uptake through both *Tg*ApiAT1 and *Tg*ApiAT6-1 modulate the Arg-dependent regulation of *Tg*ApiAT1. The loss of *Tg*ApiAT1 and *Tg*ApiAT6-1, and an increase in [Lys] in the growth medium, are all predicted to result in a depletion of cytosolic [Arg] in the parasite [15]. Our data in Figures 3 and 4 are therefore consistent with the hypothesis that the parasite is able to sense [Arg] in its cytosol, and respond to changes in cytosolic [Arg] by regulating *Tg*ApiAT1 expression.

### *T. gondii* parasites modulate *Tg*ApiAT1 expression *in vivo*

Our data to this point indicate that *T. gondii* parasites are able to sense and respond to changes in [Arg] in their environment. We hypothesise that this enables parasites to modulate Arg uptake through *Tg*ApiAT1 as they encounter different [Arg] during an infection. To investigate whether parasites vary their expression of *Tg*ApiAT1 *in vivo*, we infected mice with dual reporter strain parasites expressing nanoLUC from the wild type *Tg*ApiAT1 5’ region. Seven days after infection, we measured the nanoLUC:fLUC ratio in parasites extracted from a range of organs and from the peritoneal cavity. The ratio varied significantly between organs, with the highest ratios found in the liver, and the lowest in the spleen and kidneys (Figure 5A). The different ratios observed in parasites harvested from different organs are consistent with the parasites encountering different [Arg] in these organs during infection. Comparison of the nanoLUC:fLUC luminescence ratios in each organ to those measured in the *in vitro* experiments indicate that *T. gondii* parasites encounter an [Arg] range of ∼10-100 µM *in vivo* (Figure 5B).

**Figure 5.**
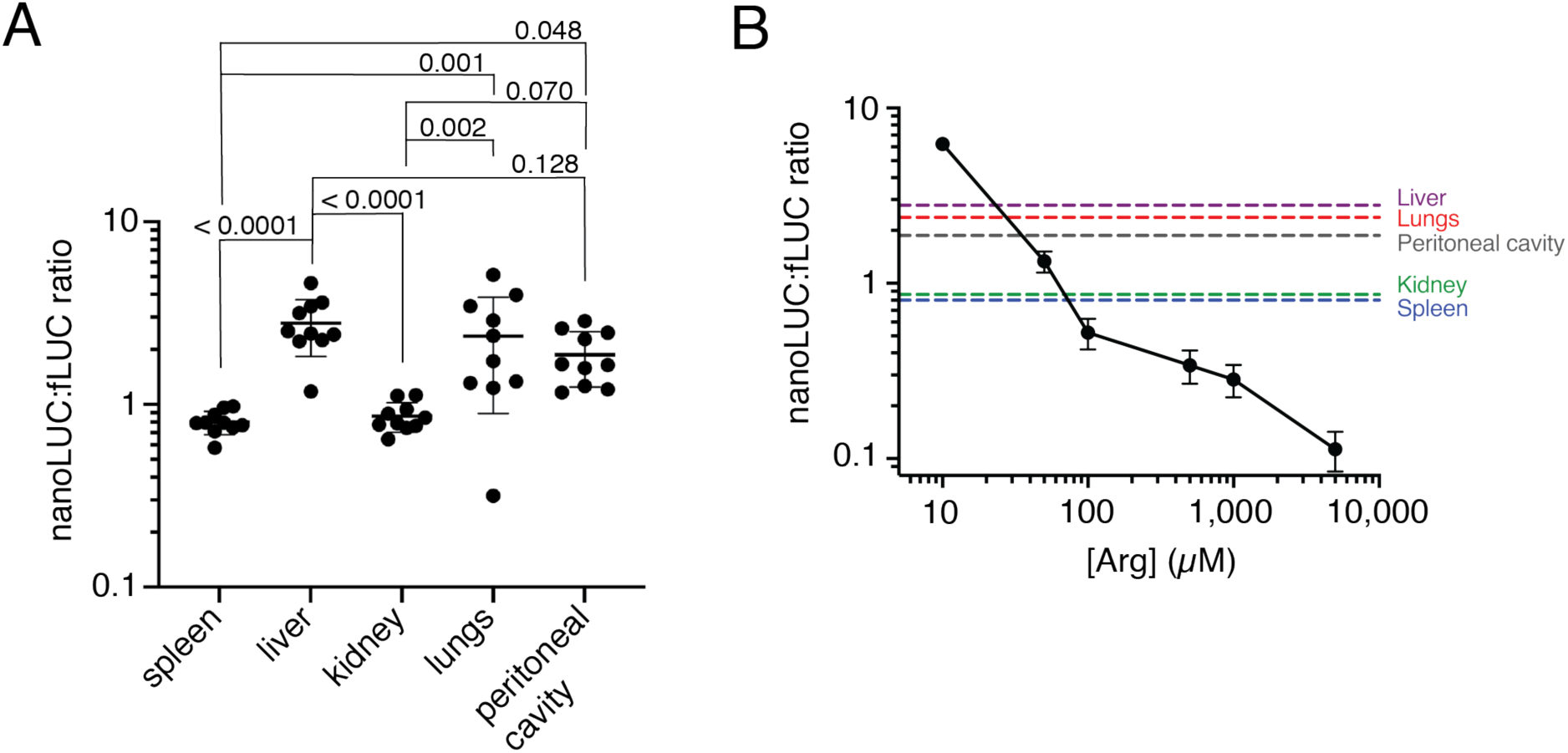
*T. gondii* parasites modulate *Tg*ApiAT1 expression *in vivo*. (**A**) NanoLUC:fLUC ratios in WT parasites harvested from a range of organs from infected mice. Mice were infected intraperitoneally with 10^3^ parasites, and euthanised seven days post-infection. Data were derived from two independent experiments with 5 mice each. *P* values were calculated using a one-way ANOVA with Tukey’s multiple comparisons test. *P* values not shown were > 0.600. (**B**) The mean nanoLUC:fLUC luminescence ratios of WT parasites harvested from various mouse organs and peritoneal cavity in (**A**) mapped onto the nanoLUC:fLUC luminescence ratios of parasites grown *in vitro* at a range of [Arg] (Figure 2B). *In vitro* data represent the mean ± s.e.m. from nine independent experiments.

### *Tg*ApiAT1 regulation is mediated by an upstream open reading frame

Finally, we investigated the mechanism by which the 5’ region of the *Tg*ApiAT1 gene regulates *Tg*ApiAT1 expression in response to varying [Arg]. The most common mechanism of 5’-mediated gene regulation in eukaryotes is through regulating transcript abundance [18]. Quantitative real time PCR measurements of *Tg*ApiAT1 transcript abundance in parasites grown at 50 µM compared to 1.15 mM Arg revealed no significant differences (Figure 6A), indicating that Arg-dependent *Tg*ApiAT1 regulation occurs post-transcriptionally.

**Figure 6.**
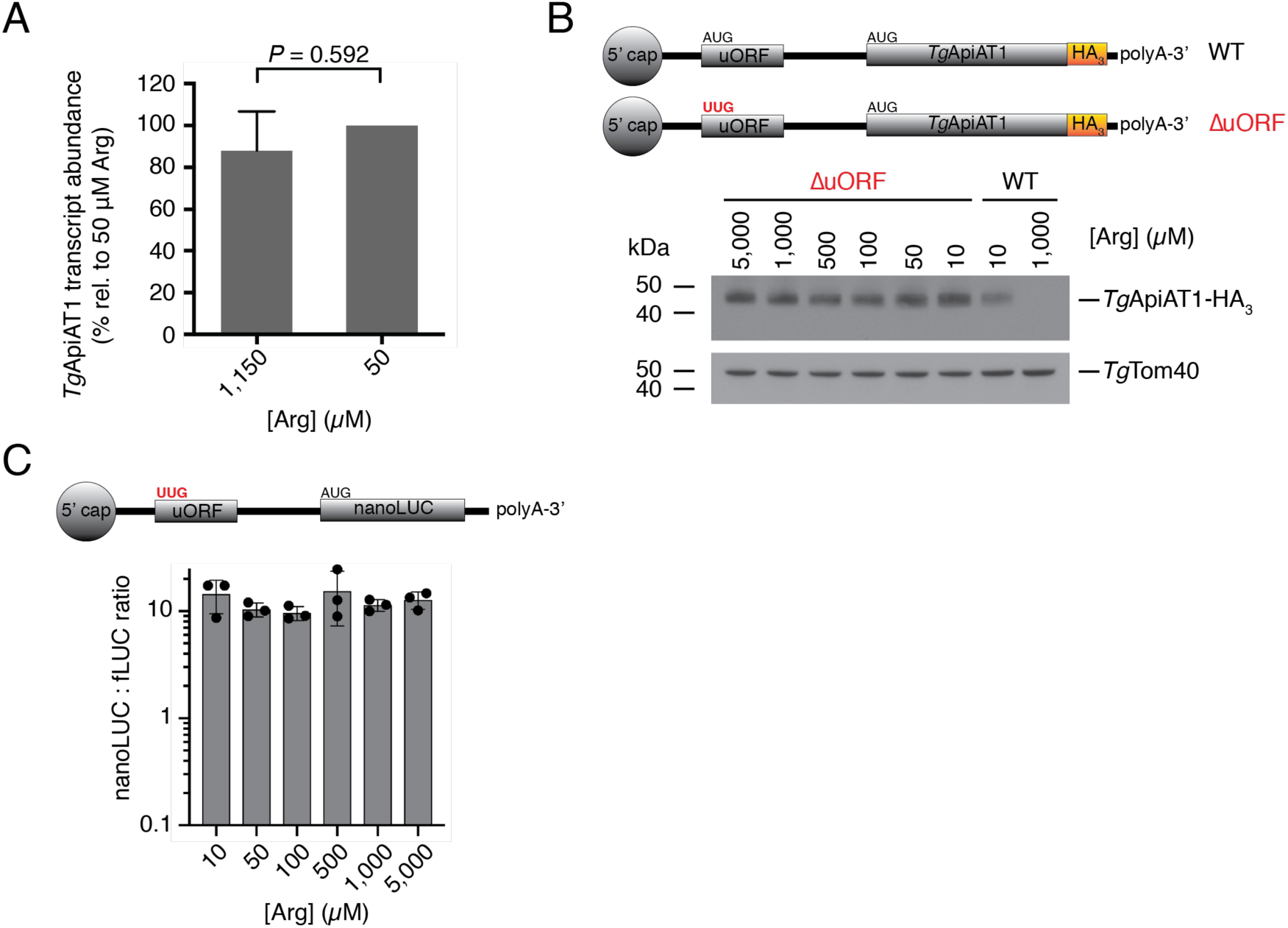
Arg-dependent regulation of *Tg*ApiAT1 occurs post-transcriptionally, and is mediated by an upstream open reading frame. (**A**) Relative *Tg*ApiAT1 transcript abundance in parasites grown at 50 µM and 1.15 mM Arg, normalised to the 50 µM condition. Data represent the mean ± SD from three independent experiments, and the *P* value was calculated using a Student’s t-test. (**B**) Western blot of ΔuORF *Tg*ApiAT1-HA_3_ parasites grown at a range of [Arg] in the growth medium, and probed with anti-HA antibodies. Western blots of WT *Tg*ApiAT1-HA_3_ parasites cultured in 10 µM or 1 mM Arg are shown for comparison. *Tg*Tom40 is a loading control. Data are representative of three independent experiments. (**C**) nanoLUC:fLUC ratio in a parasite strain expressing nanoLUC from the *Tg*ApiAT1 5’ region that lacks the uORF start codon (ΔuORF) and fLUC from the *α*-tubulin 5’ region, and grown at a range of [Arg]. Data represent the mean ± SD from three independent experiments, and were analysed using a one-way ANOVA with Tukey’s multiple comparisons test. All calculated *P* values were 0.551 or greater (not shown).

Post-transcriptional regulation can be mediated by upstream open reading frames (uORFs) in the 5’ untranslated region (5’ UTR) of transcripts [19]. We examined the *Tg*ApiAT1 5’ UTR for potential uORFs, and identified four candidate upstream ATG start codons, of which one was conserved in related the coccidian parasites *Neospora caninum* and *Sarcocystis neurona* (see below). To test whether the conserved uORF has a role in *Tg*ApiAT1 regulation, we used a CRISPR/Cas9 genome editing strategy to convert the uORF ATG to TTG in *Tg*ApiAT1-HA_3_-expressing parasites, generating a parasite strain we termed ΔuORF (Figure 6B). When this strain was exposed to varying [Arg] there was no Arg-dependent regulation of *Tg*ApiAT1-HA_3_ protein levels (Figure 6B), implicating the conserved uORF in the Arg-dependent response. We also generated a dual reporter strain in which nanoLUC was expressed from the 5’ region of *Tg*ApiAT1 lacking the uORF ATG (Figure 6C). We measured the nanoLUC:fLUC luminescence ratio in these parasites grown at a range of [Arg]. Again, we observed no significant Arg-dependent regulation of expression from the 5’ region of *Tg*ApiAT1 (Figure 6C). Together, these data indicate that Arg-dependent regulation of *Tg*ApiAT1 is uORF-mediated.

uORFs can regulate protein translation in a range of ways, including, in a few instances, by the peptide that is encoded by uORF [19, 20]. The peptide sequence encoded by the *Tg*ApiAT1 uORF peptide sequence is conserved in closely related coccidian parasites such as *N. caninum* and *S. neurona* (Figure 7A). To test whether the peptide sequence of the *Tg*ApiAT1 uORF is important for regulating translation of the downstream main ORF, we mutated the conserved aspartate residue at position 19 of the *Tg*ApiAT1 uORF to asparagine (D19N; a mutation mediated by a single base pair change in the transcript; Figure 7A) and used the mutated *Tg*ApiAT1 5’ UTR to drive nanoLUC expression in a dual reporter strain. We grew D19N parasites in media containing a range of [Arg] and measured nanoLUC:fLUC luminescence ratios. In contrast to a WT control, the nanoLUC:fLUC ratio in D19N parasites did not decrease with increasing [Arg] at most concentrations tested, although we observed a slight but significant reduction in the nanoLUC:fLUC ratio at 5 mM (Figure 7B). Expression from the *Tg*ApiAT1 5’ UTR was, therefore, largely unresponsive to variations in [Arg] in D19N parasites, consistent with the hypothesis that the peptide sequence of the *Tg*ApiAT1 uORF is important for Arg-dependent regulation.

**Figure 7.**
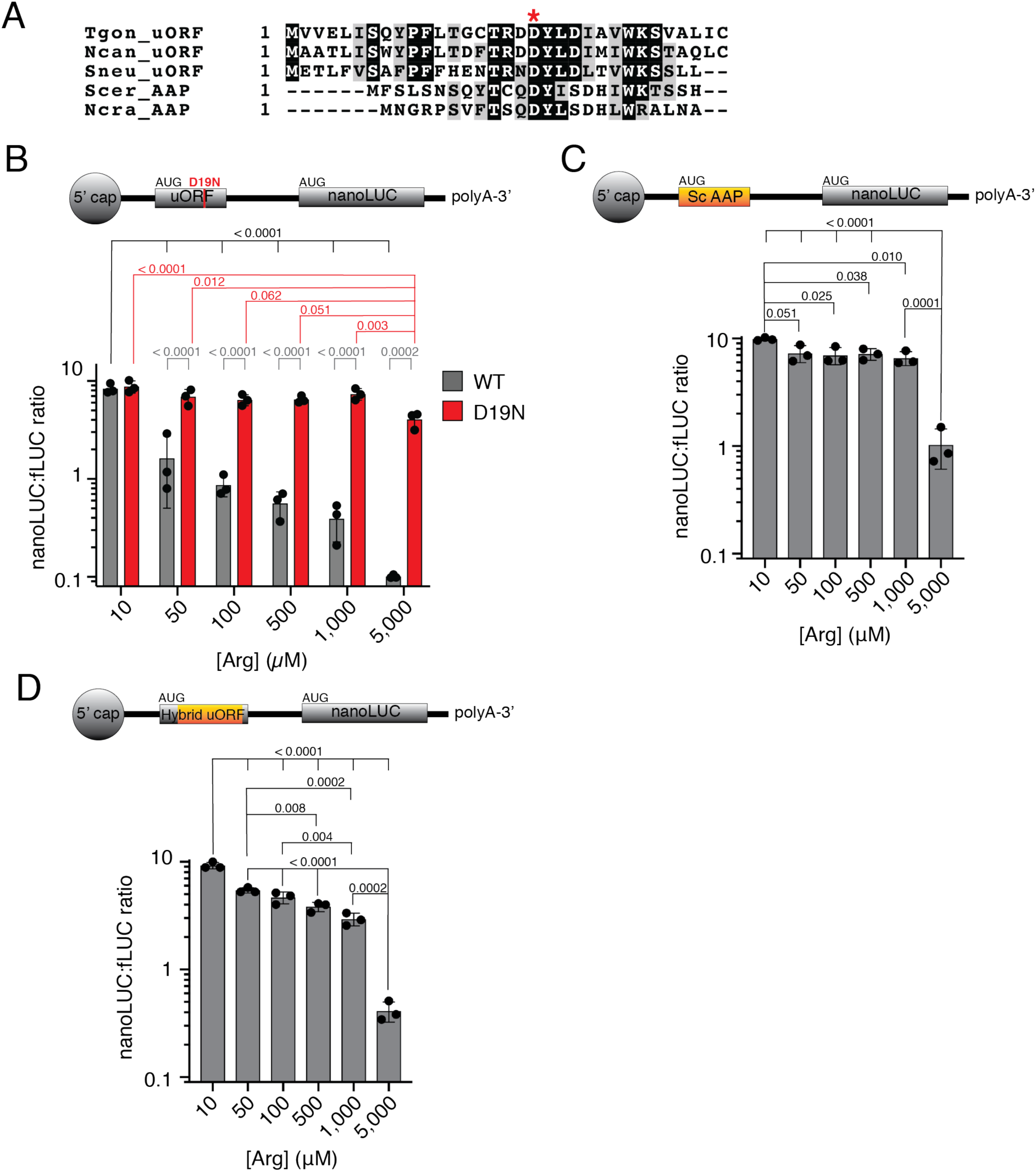
The *Tg*ApiAT1 uORF resembles the Arginine Attenuator Peptide of fungi, and mediates regulation of *Tg*ApiAT1 in a peptide sequence-dependent manner. (**A**) Multiple sequence alignment of the uORF-encoded peptide sequences of ApiAT1 homologues in *T. gondii* (Tgon_uORF) and the related coccidian parasites *Neospora caninum* (Ncan_uORF) and *Sarcocystis neurona* (Sneu_uORF), and the arginine attenuator peptides of the fungi *Saccharomyces cerevisiae* (Scer_AAP) and *Neurospora crassa* (Ncra_AAP). The conserved aspartate residue at position 19 of the *Tg*ApiAT1 uORF is highlighted with an asterisk. (**B**) NanoLUC:fLUC ratios in WT and *Tg*ApiAT1^uORF D19N^ (D19N) parasites grown at a range of [Arg]. Data represent the mean ± SD from three independent experiments. *P* values were calculated using a one-way ANOVA with Tukey’s multiple comparisons test. *P* values not shown were > 0.200. (**C-D**) NanoLUC:fLUC ratios in (**C**) *Tg*ApiAT1^ScAAP^ or (**D**) *Tg*ApiAT1^hybrid uORF^ parasites grown at a range of [Arg]. Data represent the mean ± SD from three independent experiments. *P* values were calculated using a one-way ANOVA with Tukey’s multiple comparisons test. *P* values not shown were > 0.200.

The best characterised example of uORF peptide-mediated regulation in the literature is the so-called arginine attenuator peptide (AAP) of fungi [21]. This peptide is encoded by the uORF of the gene encoding carbamoyl phosphate synthetase (Arg2), an arginine biosynthesis enzyme. Like the uORF peptide of *Tg*ApiAT1, the AAP is responsive to Arg, and mediates repression of the downstream open reading frame under arginine-replete conditions [22]. The *Tg*ApiAT1 uORF peptide has some sequence similarity to the AAP from *Saccharomyces cerevisiae* and *Neurospora crassa*, including in the conserved aspartate that is critical for both *Tg*ApiAT1 uORF and AAP function (Figure 7A-B; [21]). To test whether the *S. cerevisiae* (*Sc*AAP) can replace the function of the *Tg*ApiAT1 uORF peptide, we expressed nanoLUC from a modified *Tg*ApiAT1 5’ region in which the native uORF was replaced by a uORF encoding *Sc*AAP in a dual reporter strain. We grew the resultant strain at a range of [Arg] and measured the nanoLUC:fLUC ratio. We observed a small but significant decrease in the nanoLUC:fLUC ratio with increasing [Arg], most noticeably at the highest [Arg] tested (Figure 7C).

The *Tg*ApiAT1 uORF peptide is larger (33 amino acids) than *Sc*AAP (25 amino acids). We generated a ‘hybrid’ uORF that encoded the first seven and last two amino acids of the *Tg*ApiAT1 uORF either side of the *Sc*AAP, generating a peptide of the same length as the *Tg*ApiAT1 uORF, and incorporated this into the *Tg*ApiAT1 5’ region driving nanoLUC in a dual reporter strain. We measured the nanoLUC:fLUC ratio at a range of [Arg] and observed a significant decrease in the ratio with increased [Arg] (Figure 7D). Together, these data indicate that *Sc*AAP can partially complement the function of the *Tg*ApiAT1 uORF in mediating Arg-dependent regulation in *T. gondii*, suggesting that similar mechanisms of peptide sequence-dependent regulation may be occurring.

## DISCUSSION

This paper describes what is, to our knowledge, the first example of substrate-mediated regulation of a transporter in apicomplexan parasites. It adds to a growing body of literature on the ability of apicomplexan parasites to sense and respond to changes in nutrient availability [7–10]. Our data indicate that *T. gondii* parasites can sense [Arg] in their environment, and respond by regulating the abundance of the Arg transporter *Tg*ApiAT1. The ability of the parasite to regulate *Tg*ApiAT1 abundance may contribute to enabling the parasite to take up sufficient Arg to facilitate its proliferation as it encounters variable [Arg] across the course of an infection, and may play a role in the ability of *T. gondii* to infect a broad range of cell types in different hosts.

Arg uptake into *T. gondii* is mediated by the combined action of *Tg*ApiAT1, a selective Arg transporter, and *Tg*ApiAT6-1, a broad cationic amino acid transporter that is particularly important for Lys uptake into the parasite ([15]; Rajendran, Fairweather *et al*. unpublished). *Tg*ApiAT6-1 is constitutively expressed, regardless of the cationic amino acid concentrations that the parasites encounters (Figure 4), whereas *Tg*ApiAT1 abundance is influenced in an antagonistic manner by the concentrations of Arg and Lys in the growth medium (Figure 1; Figure 3), and is expressed at different levels in different organs during infection (Figure 5). We hypothesise that *Tg*ApiAT1 regulation enables *T. gondii* parasites to respond to different [Arg] that parasites encounter in different *in vivo* environments, enabling parasites to modulate Arg uptake from the host cell, thereby exerting tight control over their intracellular [Arg].

Regulation of cationic amino acid transporters in response to the availability of their substrates is observed in a range of organisms, including mammals and the protozoan parasite *Leishmania donovani* [23, 24]. A recent study by Augusto and colleagues demonstrated that *T. gondii*-mediated depletion of [Arg] in mammalian host cells resulted in increased abundance of the CAT1 cationic amino acid transporter of mammalian host cells [25]. Our data indicate an additional layer of complexity in mediating Arg acquisition by the parasite, with parasites able to modulate the amount of Arg that they take up from their host by regulating the level of expression of their primary Arg uptake transporter.

Arg and Lys were not the only nutrients found to regulate the abundance of *Tg*ApiAT1; *Tg*ApiAT1 abundance decreased in response to decreased levels of glucose, and the amino acids glutamine and tyrosine (Figure 3). This occurred at concentrations of these nutrients that are close to, or below, the levels required for optimal parasite growth [14, 17]. Thus, in addition to being regulated by an Arg-specific mechanism, *Tg*ApiAT1 abundance may be regulated as part of a more general starvation response. The response of *T. gondii* parasites to glutamine starvation has some similarities to the GCN2-dependent translational regulation that occurs during the starvation response of mammalian cells [17]. The putative starvation response was observed both when measuring *Tg*ApiAT1 protein abundance, and in a strain in which nanoLUC was driven by the 5’ region of the *Tg*ApiAT1 gene. This is consistent with the starvation response being mediated by the 5’ region of *Tg*ApiAT1. It remains to be determined whether this process is translationally mediated, and whether there is functional overlap between this general starvation response and the specific Arg-dependent response.

Our data indicate that parasites vary the expression of *Tg*ApiAT1 in different organs during a mouse infection (Figure 5), which may reflect differences in host Arg metabolism and, consequently, [Arg] in these organs. Several host cell enzymes catalyse reactions for which Arg is a substrate. The enzyme arginase catalyses the conversion of Arg to ornithine. Arginase activity is particularly high in the liver of mammals [26], which may explain why parasites encounter low levels of Arg in this organ. Parasites may also encounter different [Arg] across the course of an infection. Host cell nitric oxide synthases, including endothelial nitric oxide synthase (eNOS) and inducible nitric oxide synthase (iNOS), catalyse the conversion of Arg to nitric oxide (NO). eNOS is expressed in a range of cell types, and is upregulated in response to *T. gondii* infection [27], and iNOS is upregulated in an interferon *γ*-mediated innate immune response that occurs upon *T. gondii* infection [28]. Having established a means of estimating the [Arg] that parasites encounter *in vivo* (Figure 5), it will now be of interest to determine whether *Tg*ApiAT1 expression changes across the course of an infection in response to the upregulation of Arg-dependent enzymes such as eNOS and iNOS, and whether *Tg*ApiAT1 regulation plays a role in enabling parasite proliferation and dissemination to different organs and tissues as an infection progresses.

We demonstrated that Arg-dependent regulation of *Tg*ApiAT1 is mediated by a uORF in the *Tg*ApiAT transcript (Figures 6-7). uORFs appear to be abundant in *T. gondii* transcripts [29], and the uORF of *Tg*ApiAT1 represents the first characterised example of a functional uORF in these parasites. Our data indicate that the peptide encoded by the uORF plays a role in the Arg-dependent regulation of *Tg*ApiAT1 expression (Figure 7). This is one of only a few known cases in which the peptide of a uORF appears to be critical for regulating translation of the downstream main ORF [19]. The best studied example of peptide-dependent uORF regulation is the AAP of fungi, which regulates the Arg-dependent translation of the arginine biosynthesis enzyme Arg2 [30]. The AAP mediates ribosome stalling on the Arg2 transcript in Arg-replete conditions, possibly by blocking the ribosome exit tunnel [22, 31, 32]. The sequence of the *Tg*ApiAT1 uORF resembles that of the AAP (Figure 7A), with one of the conserved residues being critical for *Tg*ApiAT1 uORF function, and the yeast AAP being partially functional in *T. gondii* (Figure 7B-D). Given that *T. gondii* and fungi are separated by ∼1.5 billion years of evolution, and that the *Tg*ApiAT1 uORF peptide appears restricted to *T. gondii* and its closest relatives, a conserved function between these uORFs would represent a remarkable example of convergent evolution.

## Methods

### Parasite culture

Parasite cultures were maintained in human foreskin fibroblasts (a kind gift from Holger Schlüter, Peter MacCallum Cancer Centre) in a humidified 37°C incubator at 5 % CO_2_. Host cells were checked periodically for *Mycoplasma* infection. Unless otherwise indicated in the text, parasites were cultured in RPMI supplemented with 1 % (v/v) foetal calf serum, 2 mM glutamine, 50 U/ml penicillin, 50 µg/ml streptomycin, 10 µg/ml gentamicin, and 0.25 µg/ml amphotericin b, as described [15]. For all ‘homemade’ media where we varied the concentrations of nutrients, we used 1% (v/v) dialysed foetal calf serum. Where applicable, ATc was added to a final concentration of 0.5 µg/ml. Experiments to measure the effects of a range of [Arg] on *Tg*ApiAT1-HA_3_ abundance or *apiAT1* 5’-nanoLUC activity were performed in RPMI containing 200 µM Lys, unless otherwise indicated. Experiments to measure the effects of [Lys], [Tyr], [Gln] and [D-glucose] on *Tg*ApiAT1 regulation were performed in medium containing 50 µM Arg. Plaque assays were performed in 25 cm^2^ tissue culture flasks, with 500 parasites added to a flask. Parasites were grown for 8 days before being stained in a solution of 2 % (w/v) crystal violet, 20 % (w/v) ethanol and 0.8 % (w/v) ammonium acetate. To induce bradyzoite formation, 1.4 × 10^6^ tachyzoites were inoculated into a 25 cm^2^ tissue culture flask with confluent human foreskin fibroblasts and allowed to proliferate for 20 hr in standard growth medium. The growth medium was replaced with alkaline RPMI supplemented with 25 mM HEPES (pH 8.2-8.4), and the infected host cells were then cultured for a further six days at ambient CO_2_ levels. Intracellular bradyzoites were mechanically egressed from host cells using a 26 gauge needle, then further disrupted using a 30 gauge needle before sample preparation for SDS-PAGE.

### Ethics Statement

All animal research was conducted in accordance with the National Health and Medical Research Council’s Australian Code for the Care and Use of Animals for Scientific Purposes, and the Australian Capital Territory Animal Welfare Act 1992. Mice were maintained and handled in accordance with protocols approved by the Australian National University Animal Experimentation Ethics Committee (protocol number A2016/42).

### Mouse infections

Freshly egressed, dual reporter strain parasites were filtered through a 3 µm polycarbonate filter, washed once in phosphate-buffered saline (PBS), and resuspended to 1 × 10^4^ parasites/ml in PBS. 6-8 week-old, female Balb/c mice were inoculated intraperitoneally with 1 × 10^3^ parasites using a 26-gauge needle. Mice were weighed regularly and monitored for symptoms of toxoplasmosis (weight loss, ruffled fur, lethargy and hunched posture). At day 6, mice were imaged using an IVIS imaging system to confirm infection, as described [33]. Briefly, mice were injected intraperitoneally with 200 µl of 15 mg/ml D-luciferin in PBS, anaesthetised with 2.5 % isofluorane in oxygen in an anaesthetic chamber using an XGI-8 anaesthesia system, and imaging was performed on an IVIS Spectrum imaging system 10 min post-injection. Anaesthesia was maintained during imaging by application of 2.5% isofluorane in oxygen via a nose cone. All mice were euthanised at day 7 of the experiment and dissected to remove organs for dual luciferase assay measurements, as described below. We also euthanised and analysed two uninfected mice to determine background luminescence levels found in each tested organ.

### Generation of genetically modified *T. gondii* strains

To incorporate a 3xHA tag into the *Tg*ApiAT1 locus, we adopted a CRISPR/Cas9 genome editing strategy. We introduced a single guide RNA (gRNA) targeting the 3’ region of the *Tg*ApiAT1 locus into the vector pSAG1::Cas9-U6::sgUPRT (Addgene plasmid # 54467; [34]) using Q5 site-directed mutagenesis (New England Biolabs) with the primers ApiAT1 3’ gRNA fwd and generic rvs (Table S2), as described previously [34]. We generated a donor DNA containing the 3xHA-tag flanked by sequence homologous to the *Tg*ApiAT1 locus either side of the *Tg*ApiAT1 stop codon as a gBlock (IDT; Table S2). We amplified the ‘*Tg*ApiAT1-HA_3_’ gBlock DNA (IDT) by polymerase chain reaction (PCR) using the primers ApiAT1 3’ edit fwd and rvs (Table S2). We co-transfected the gRNA/Cas9-GFP-expressing vector and the donor DNA into TATiΔ*ku80* [35], PrugniaudΔ*ku80Δhxgprt*/*ldh2*-GFP [36], or r*Tg*ApiAT6-1 strain parasites, and sorted GFP-expressing clones 2-3 days post-transfection, as described [14, 37].

To generate the dual luciferase reporter strain, we first generated a strain that expressed firefly luciferase (fLUC) under the control of the *T. gondii α*-tubulin 5’ region. We digested the vector pTub8-rsLUC (a kind gift from Boris Striepen, U. Penn) with *Spe*I and *Not*I and ligated this into the equivalent sites of pDTG [38], a vector that encodes a pyrimethamine-resistance marker. We transfected this plasmid into RHΔ*hxgprt* [39] strain parasites, selected on pyrimethamine, and obtained clonal parasites by limiting dilution. This generated a strain that we termed the *α*tub 5’-fLUC strain, which constitutively expressed fLUC from the *α*-tubulin 5’ region. We next set about generating a plasmid that expressed nanoLUC from the *Tg*ApiAT1 5’ region. First, we generated a vector expressing firefly luciferase (fLUC) under the control of the *Tg*ApiAT1 5’ region. We amplified fLUC with the primers fLUC fwd and fLUC rvs (Table S2) using the LT-3 plasmid [40] (a kind gift from Alex Maier, ANU) as template. We digested the resulting product with *Bgl*II and *Avr*II and ligated this into the equivalent sites of the vector pUgCTH_3_ [15], generating a vector we termed pUgCT-fLUC-HA_3_. We PCR amplified the 1.2 kb region upstream of the *Tg*ApiAT1 5’ UTR (*i.e.* upstream of the transcript start site) using the primers ApiAT1 5’ fwd and rvs (Table S2), digested the product with *Spe*I and *Asi*SI and ligated into the equivalent sites of pUgCT-fLUC-HA_3_. We then amplified the 5’ UTR of the *Tg*ApiAT1 gene using the ApiAT1 5’ UTR fwd and rvs primers (Table S2), digested the resulting product with *Sbf*I and *Asi*SI, and ligated this into the equivalent sites of the pUgCT-fLUC-HA_3_ vector, terming the resultant vector pUgC-apiAT1 5’-fLUC-HA_3_. Next, we amplified nanoLUC using the nanoLUC fwd and rvs primers (Table S2) and the plasmid pTubNluc-AID-2xHA-DHFR (a kind gift from Boris Striepen, U. Penn) as template. We digested the resulting product with *Asi*SI and *Avr*II, and ligated this into the equivalent site of pUgC-apiAT1 5’-fLUC-HA_3_, generating a vector we termed pUgC-apiAT1 5’-nanoLUC-HA_3_. The resultant plasmid encodes nanoLUC under control of the *Tg*ApiAT1 5’ region. We transfected this plasmid into the *α*tub 5’-fLUC parasite strain, selected on chloramphenicol, and obtained clonal parasites by limiting dilution. We termed the resultant strain the ‘dual reporter strain’.

To generate a *T. gondii* strain in which we could knock down expression of *Tg*ApiAT6-1, we replaced the native *Tg*ApiAT6-1 promoter region with an ATc-regulatable promoter using a double homologous recombination approach. First, we amplified the 5’ flank of *Tg*ApiAT6-1 with the primers ApiAT6-1 5’ flank fwd and rvs (Table S2). We digested the resulting product with *Psp*OMI and *Nde*I and ligated into the equivalent sites of the vector pPR2-HA_3_ [41], generating a vector we termed pPR2-HA_3_(ApiAT6-1 5’ flank). Next, we amplified the 3’ flank with the primers ApiAT6-1 3’ flank fwd and rvs (Table S2). We digested the resulting product with *Bgl*II and *Not*I and ligated into the equivalent sites of the vector pPR2-HA_3_(ApiAT6-1 5’ flank) vector. We linearised the resulting plasmid with *Not*I and transfected this into TATi/Δ*ku80* strain parasites [35] expressing tandem dimeric Tomato RFP. We selected, on pyrimethamine, and cloned parasites by limiting dilution. We termed the resulting strain regulatable (r)*Tg*ApiAT6-1. To enable us to measure knockdown of the ATc-regulatable *Tg*ApiAT6-1 protein, we integrated a HA tag into the r*Tg*ApiAT6-1 locus by transfecting a *Tg*ApiAT6-1-HA 3’ replacement vector, described previously [14], into this strain. To incorporate a 3xHA tag into the *Tg*ApiAT1 locus of the r*Tg*ApiAT6-1 strain, we adopted a CRISPR/Cas9 genome editing strategy, as described above.

To generate a frameshifted ‘knockout’ mutation in the *Tg*ApiAT1 locus of the dual reporter strain, we transfected this with a plasmid expressing a gRNA targeting the *Tg*ApiAT1 locus, sorted and cloned parasites 3 days after transfection, and verified that a successful frameshift mutation (a single base pair insertion) had occurred by sequencing the *Tg*ApiAT1 locus, all as described previously [14].

To generate a *Tg*ApiAT1-HA_3_-expressing strain wherein the ATG start codon of the *Tg*ApiAT1 uORF was mutated to TTG, we adopted a CRISPR/Cas9 genome editing strategy. First, we introduced a gRNA targeting the genomic locus that encoded the *Tg*ApiAT1 5’ UTR near the uORF start codon into pSAG1::Cas9-U6::sgUPRT vector using Q5 site-directed mutagenesis with the primers ApiAT1 uORF gRNA fwd and generic rvs (Table S2) as described previously [34]. We generated a donor DNA wherein the ATG of the uORF was mutated to TTG by annealing the complementary primers ApiAT1 ΔuORF fwd and rvs (Table S2), and co-transfected this with the gRNA-expressing vector into *Tg*ApiAT1-HA_3_ strain parasites. We sorted GFP-expressing clones 3 days post-transfection, then sequenced clones to verify successful mutation. In addition to the ATG start codon of the uORF being mutated to TTG, the clone that we characterised had an additional G to C mutation in the protospacer adjacent motif (PAM) site of the gRNA target (13 bp upstream of the ATG codon) designed to prevent gRNA-mediated Cas9 cutting the chromosome following genome modification, and an unintended G to A mutation 6 bp upstream of the start codon, likely introduced by a mutation in the donor DNA.

To generate a strain expressing nanoLUC from the *Tg*ApiAT1 5’ region in which the uORF ATG start codon was mutated to TTG, we amplified the 5’UTR of the *Tg*ApiAT1 using the ApiAT1 5’ UTR fwd and rvs primers (Table S2), and a ‘*Tg*ApiAT1/ΔuORF 5’UTR’ gBlock (IDT) encoding an altered *Tg*ApiAT1 5’ UTR region in which the start codon of the *Tg*ApiAT1 uORF was mutated to TTG (Table S2). We digested the resultant PCR product with *Pst*I and *Asi*SI, and ligated into the *Sbf*I and *Asi*SI sites of pUgC-apiAT1 5’-nanoLUC-HA_3_. We transfected this vector into the *α*tub 5’-fLUC strain, selected on chloramphenicol, and obtained clonal parasites by limiting dilution. To generate a strain expressing nanoLUC from the *Tg*ApiAT1 5’ region wherein the native *Tg*ApiAT1 uORF was replaced with the *S. cerevisiae* AAP uORF, we amplified a modified *Tg*ApiAT1 5’UTR containing the *S. cerevisiae* AAP uORF using the ApiAT1 5’ UTR fwd and rvs primers (Table S2) and a ‘*Tg*ApiAT1/ScAAP uORF 5’UTR’ gBlock (IDT; Table S2). We digested the resultant PCR product with *Pst*I and *Asi*SI, and ligated into the *Sbf*I and *Asi*SI sites of pUgC-apiAT1 5’-nanoLUC-HA_3_. We transfected this vector into the *α*tub 5’-fLUC strain, selected on chloramphenicol, and obtained clonal parasites by limiting dilution. To generate a strain expressing nanoLUC from the *Tg*ApiAT1 5’ region containing a ‘hybrid’ uORF consisting of the *S. cerevisiae* AAP flanked by the 5’ and 3’ regions of the *Tg*ApiAT1 uORF, we amplified a modified *Tg*ApiAT1 5’UTR containing the hybrid *Tg*ApiAT1 uORF using the ApiAT1 5’ UTR fwd and rvs primers (Table S2) and a ‘*Tg*ApiAT1/hybrid uORF 5’UTR’ gBlock (IDT; Table S2). We digested the resultant PCR product with *Pst*I and *Asi*SI, and ligated into the *Sbf*I and *Asi*SI sites of pUgC-apiAT1 5’-nanoLUC-HA_3_. We transfected this vector into the *α*tub 5’-fLUC strain, selected on chloramphenicol, and obtained clonal parasites by limiting dilution.

To generate a strain expressing nanoLUC from the *Tg*ApiAT1 5’ region wherein the aspartate residue at position 19 of the uORF peptide was mutated to asparagine (D19N), we used Q5 mutagenesis approach. We followed the manufacturer’s instructions (New England Biolabs) using the pUgC-apiAT1 5’-nanoLUC-HA_3_ plasmid as template, and the uORF D19 fwd and rvs primers (Table S2). We transfected the resultant vector into *α*tub 5’-fLUC strain parasites, selected on chloramphenicol, and obtained clonal parasites by limiting dilution.

To generate a strain that expressed nanoLUC from the *α*-tubulin 5’ region, we amplified the *α*-tubulin 5’ region with the primers Tub 5’ fwd and rvs (Table S2), digested the product with *Spe*I and *Asi*SI and ligated into the equivalent sites of the pUgC-apiAT1 5’-nanoLUC-HA_3_, generating a vector we termed pUgC-tub 5’-nanoLUC-HA_3_. We transfected this plasmid into RHΔ*hxpgrt* strain parasites, selected on chloramphenicol, and obtained clonal parasites by limiting dilution.

### Quantitative real time PCR

TATiΔ*ku80* strain parasites were cultured for 2 days in modified RPMI medium containing 50 µM or 1.15 mM Arg. Parasites were mechanically egressed from host cells using a 26 gauge needle, then total RNA was extracted using the Isolate II RNA mini extraction kit (Bioline), according to the ‘cultured cells and tissue’ protocol in the manufacturer’s instructions. cDNA synthesis was performed using the High-Capacity cDNA reverse transcriptase kit (Applied Biosystems) with a random primer mix and 2 µg total RNA from each sample, according to the manufacturer’s instructions. Quantitative real time PCR was performed using a LightCycler 480 system (Roche) with the LightCycler 480 SybrGreen I Master mix, following the manufacturer’s instructions, and using 5 µM primers. The LightCycler 480 conditions were as follows: 10 min preincubation at 95°C, then 45 cycles of 15 sec denaturation (95°C), 15 sec annealing (58°C), and 20 sec elongation (72°C). To detect the abundance of *Tg*ApiAT1 transcript, we used the primers ApiAT1 qrt int fwd and rvs (which amplified *Tg*ApiAT1 cDNA across the intron of the transcript) and ApiAT1 qrt 3’ UTR fwd and rvs (which amplified *Tg*ApiAT1 cDNA from the 3’ UTR of the transcript; Table S2). *Tg*ApiAT1 transcript levels were normalised using *α*-tubulin (Tub) and glyceraldehyde-3-phosphate dehydrogenase (GAPDH) as housekeeping transcript controls. We amplified these housekeeping controls using the primers Tub qrt fwd and rvs and GAPDH qrt fwd and rvs; Table S2). Raw fluorescence data were exported and analysed using LinRegPCR[42] to perform background subtraction and determine PCR primer efficiency. Samples were then normalized to housekeeping controls using the Pfaffl equation (E_NPT1_^ΔCTNPT1^/E_ref_^ΔCTref^; E=primer efficiency, ΔCT = difference in cycle threshold between samples grown at 1.15 mM and 50 µM Arg, ref = housekeeping controls; [43]) and expressed as percentage relative to that at 50 µM. Three biological replicates were performed and each reaction was done in at least triplicate.

### Western blotting

Protein samples were separated using NuPAGE Bis/Tris gels, as described [15], loading 2.5 × 10^6^ parasite equivalents per lane. Membranes were probed with rat anti-HA antibodies (1:100 to 1:3,000 dilutions; clone 3F10, Sigma, 11867423001), rabbit anti-*Tg*Tom40 [44] (1:2,000 dilution), mouse anti-GFP (1:1,000 dilution; Sigma, 11814460001), mouse anti-BAG1 [45] (1:250 dilution; a kind gift from Louis Weiss, Albert Einstein College of Medicine), rabbit anti-SAG1 (1:1,000 dilution; a kind gift from Michael Panas and John Boothroyd, Stanford University), or mouse anti-*Tg*GRA8 [46] (1:100,000 dilution; a kind gift from Gary Ward, U. Vermont) as primary antibodies, and horseradish peroxidase-conjugated goat anti-rat (1:5,000 to 1:10,000 dilutions; Santa Cruz, sc-2006, or Abcam, ab97057), goat anti-rabbit (1:5,000 to 1:10,000 dilution; Santa Cruz, sc-2004, or Abcam, ab97051), or goat anti-mouse (1:5,000 to 1:10,000 dilution; Santa Cruz, sc-2005) secondary antibodies.

### SWATH-MS proteomic analysis

#### Sample preparation

We undertook a SWATH-MS-based quantitative proteomic approach [16] to establish whether the abundance of proteins changed in parasites grown in media containing low vs high [Arg]. We cultured RHΔ*hxgprt* strain parasites in modified DME containing 50 µM or 1.15 mM Arg, and a constant 800 µM Lys for two days. Our previous data indicate that 50 µM is the minimum [Arg] required for optimal parasite growth [15], and we chose 50 µM as the low [Arg] value (and not a lower concentration) to avoid identifying proteins that change abundance as a result of a general starvation response. We performed five replicates for each condition. Parasites were mechanically egressed through a 26 gauge needle, filtered through a 3 µm polycarbonate filter to remove host cell debris, washed in PBS, then resuspended in a lysis buffer containing 1% (w/v) sodium dodecyl sulfate (SDS), 1 mM dithiothreitol (DTT), 50 mM Tris-HCl, pH 8. SDS was removed by buffer exchange with 100 mM triethylammonium biocarbonate.

#### Sample processing

100 µg of protein from each sample was reduced in 10 mM DTT, alkylated with 20 mM iodoacetamide, then digested by trypsin for 16 hr at 37°C. Digested samples were cleaned up using a detergent removal spin column (Pierce), then dried and resuspended in 100 µL of 2% (v/v) acetonitrile with 0.1 % (v/v) formic acid. For one-dimensional information dependent acquisition (1D-IDA), 10 µl of each sample was subjected to nanoLC MS/MS analysis using an Ultra nanoLC (Eksigent) system and Triple TOP 5600 mass spectrometer (AB Sciex). For two-dimensional (2D)-IDA, a pool was prepared from 20 µl of each sample, and separated by high pH reverse phase fractionation on a Agilent 1260 quaternary HPLC system with a Zorbax 300Extend-C18 column, with 12 fractions collected. Each 1D-IDA and 2D-IDA sample was injected onto a Captrap peptide trap (Bruker) for pre-concentration and desalting in 2% (v/v) acetonitrile with 0.1 % (v/v) formic acid, then injected into the analytical column. The reverse phase nanoLC eluent was subjected to positive ion nanoflow electrospray analysis in an IDA mode. For data independent acquisition (SWATH), 10 µl of each sample was treated as for the IDA samples, with the reverse phase nanoLC eluent subjected to positive ion nanoflow electrospray in a data independent acquisition mode. For SWATH-MS, m/z window sizes were determined based on precursor m/z frequencies (m/z 400-1250) from the IDA data. In SWATH mode, a TOFMS survey scan was acquired (m/z 350-1,500, 0.05 sec) then 60 predefined m/z ranges were sequentially subjected to MS/MS analysis. MS/MS spectra were accumulated for 60 ms in the mass range 350-1,500 with optimised rolling collision energy.

#### Data processing and analysis

LC-MS/MS data from the IDA experiments were searched using ProteinPilot (v4.2; AB Sciex) against the ToxoDB GT1 proteome (ToxoDB.org). SWATH data were extracted using PeakView (v2.1) with the following parameters: the six most intense fragments of each peptide were extracted from the SWATH data sets, with shared and modified peptides excluded. Peptides with confidence ≥ 99% and FDR ≤ 1% were used for quantitation. SWATH protein peak areas were analysed using an in-house Australian Proteome Analysis Facility (APAF) program. Protein peaks were normalised to total peak area for each run, and were subjected to statistical analysis to compare relative protein peak areas between the sample groups. The data for each identified protein is presented in Table S1.

### Dual luciferase reporter assays

To measure nanoLUC and fLUC activity in dual luciferase reporter strains, we cultured parasites in 25 cm^2^ tissue culture flasks in the required growth medium. Before parasite inoculation, host cells and parasites were both were washed twice with PBS to remove residual media. Parasites were cultured for between 38 and 42 hr, over which time all remained intracellular. Parasites grown in 10 µM Arg exhibited slower growth than at other [Arg] across this timeframe. To compensate for this, we inoculated more parasites into flasks containing 10 µM Arg. On the day of the experiment, parasites were liberated from host cells by passage through a 26 gauge needle. Host cell debris were removed by filtering through a 3 µm polycarbonate filter, and parasites were pelleted by centrifugation at 1,500 × *g* for 10 min. Parasites were resuspended to 1-2 × 10^7^ parasites/ml in PBS and 25 µl of parasite suspension was added to wells of an OptiPlate-96 opaque, white 96-well plate (PerkinElmer). To measure nanoLUC and fLUC luminescence, we used the NanoGlo Dual-luciferase reporter assays system (Promega, N1610). First, we measured fLUC activity by adding 25 µl ONE-Glo Ex Luciferase assay buffer with added substrate to wells containing parasites, incubating for 5 min, then reading on a FluoStar Optima plate reader (BMG Labtech) using the luminescence settings without an emission filter. Next, we measured nanoLUC activity by adding 25 µl NanoDLR Stop & Go assay buffer containing 1:100 diluted substrate to the parasite suspension, incubating for 5 min, then reading luminescence using the same settings as for fLUC. In each assay, we included a ‘no parasite’ control (25 µl PBS), which was subtracted from the luminescence readings of the parasite-containing wells before subsequent data analysis. To measure nanoLUC and fLUC activities in mouse organs, infected and uninfected mice were euthanised by cervical dislocation. Prior to organ harvest, intraperitoneal lavage was performed by injecting 5 ml ice-cold PBS into the intraperitoneal cavity using a 26 G needle, mixing peritoneal cavity content and subsequent aspiration of the content using 20 G needle. Next, incisions were made to open the chest cavity without damaging any organs. The spleen, liver and kidneys were harvested and placed in 2 ml ice-cold PBS. Next, the lungs were perfused by injection of 10 ml of ice-cold PBS into mouse heart ventricles. The heart, lung and brain were subsequently harvested and kept in 2 ml of ice-cold PBS. All samples were kept on ice until luminescence measurements. For luminescence measurements, all organs were homogenised using a dounce homogeniser. 25µl aliquots of each of the crude homogenate samples were transferred in duplicate into wells of an OptiPlate-96 opaque, white 96-well plate. NanoLUC and fLUC measurements were performed as described above. Luminescence measurements in the heart and brain of infected mice were found to be at background levels, and were not analysed further.

### Arg uptake experiments

Experiments to measure uptake of [^14^C]Arg through *Tg*ApiAT1 were performed as described previously [47]. Briefly, extracellular *T. gondii* parasites were incubated in PBS containing 10 mM D-glucose, 0.1 µCi/ml [^14^C]Arg, 50 µM unlabelled Arg, and 800 µM unlabelled Lys for a range of times. The unlabelled Lys was added to inhibit Arg uptake through *Tg*ApiAT6-1. The reaction was stopped by centrifuging the parasites through an oil mix consisting of 84% (v/v) PM125 silicone fluid and 16 % (v/v) light mineral oil. The incorporated radiolabel was measured using a liquid scintillation counter (Perkin Elmer). Time course data were fitted by a single exponential function and the initial rate calculated from the initial slope of the curve.

### Statistics and reproducibility

Unless described otherwise in the figure legends, all quantitative data are presented as mean ± SD of three or more independent experiments. All non-quantitative data (western blots, plaque assays) displayed are representative images of multiple independent experiments, with the number of experiments listed in the figure legends. Graphs were plotted using GraphPad Prism, and statistics were also undertaken in GraphPad Prism. Details of statistics are provided in the figure legends.

## Acknowledgements

We are grateful to the students of the 2014 Biology of Parasitism Course (Marine Biological Laboratory, Woods Hole, MA) for first uncovering the Arg-mediated regulation of *Tg*ApiAT1, and students from the 2015 course for contributing further experimental insights into this process. We thank Gary Ward, Louis Weiss, Michael Panas, John Boothroyd, Alex Maier and Boris Striepen for sharing reagents, Cathy Gillespie for assistance with *in vivo* luminescence mouse imaging, and Harpreet Vohra and Michael Devoy for performing cell sorting. This work was supported by a Discovery Grant from the Australian Research Council to K.K. and G.v.D. (DP150102883) and a Project Grant from the Australian National Health and Medical Research Council (GNT1128911) to N.S. SWATH-MS proteomics was undertaken at the Australian Proteome Analysis Facility, with the infrastructure provided by the Australian Government through the National Collaborative Research Infrastructure Strategy (NCRIS)

---

**Figure S1.**
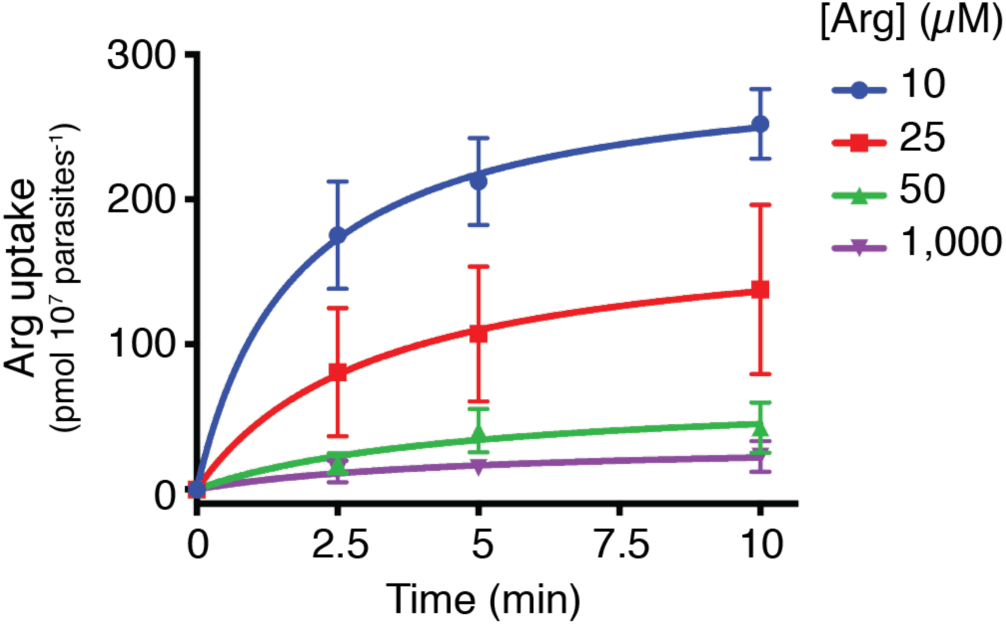
Timecourse of Arg uptake in parasites grown in medium containing a range of [Arg]. Uptake of Arg uptake in parasites cultured in growth medium containing 10, 25, 50 and 1,000 µM Arg over 10 min. Uptake was measured in 50 µM unlabelled Arg and 0.1 µCi/ml [^14^C]Arg. Data represent the mean ± s.e.m. from three independent experiments.

**Figure S2.**
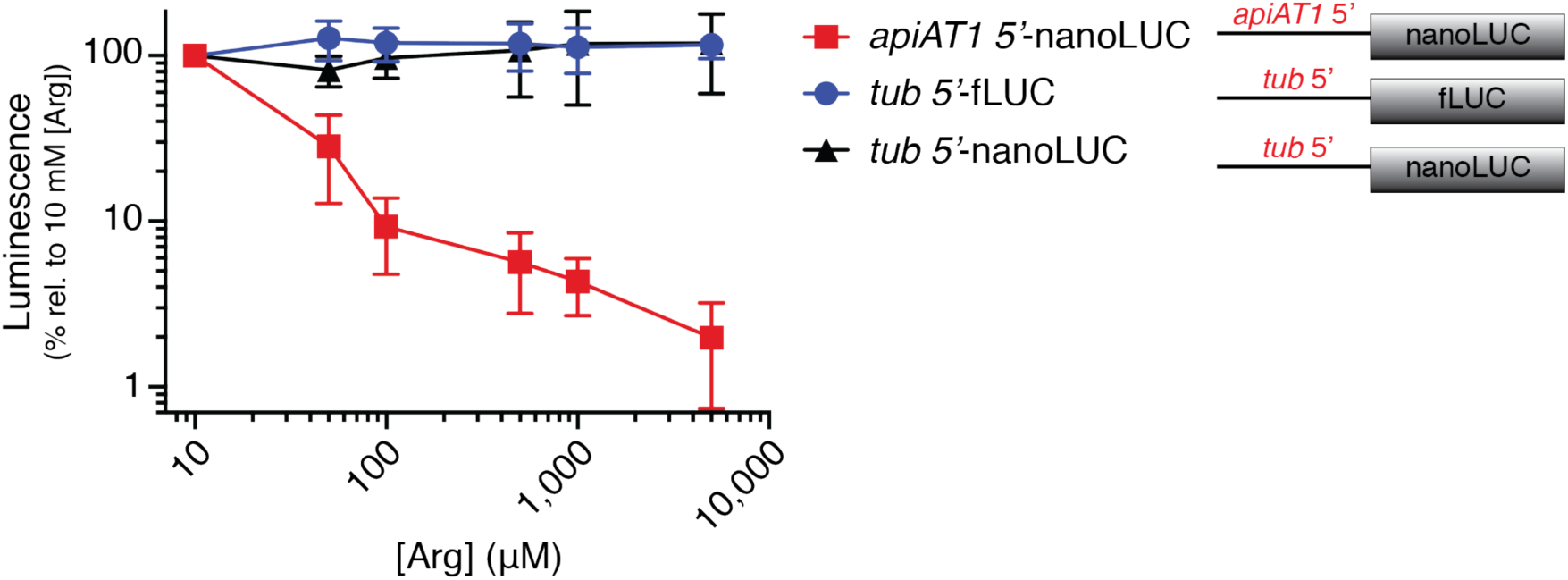
Luminescence readings of nanoLUC and fLUC expressing parasites grown at a range of [Arg]. NanoLUC and fLUC luminescence in a parasite strain expressing nanoLUC from the *Tg*ApiAT1 5’ region (*apiAT1* 5’-nanoLUC; red) and fLUC from the *α*-tubulin 5’ region (*tub* 5’-fLUC; blue), or a strain expressing nanoLUC from the *α*-tubulin 5’ region (*tub* 5’-nanoLUC; black), grown at a range of [Arg]. Luminescence is expressed as a percent of the luminescence at the 10 µM Arg condition for both nanoLUC and fLUC measurements. Data points represent the mean ± SD of nine independent experiments in the *apiAT1* 5’-nanoLUC/ *tub* 5’-fLUC strain, and the mean ± SD of four independent experiments in the *tub* 5’-nanoLUC strain.

**Figure S3.**
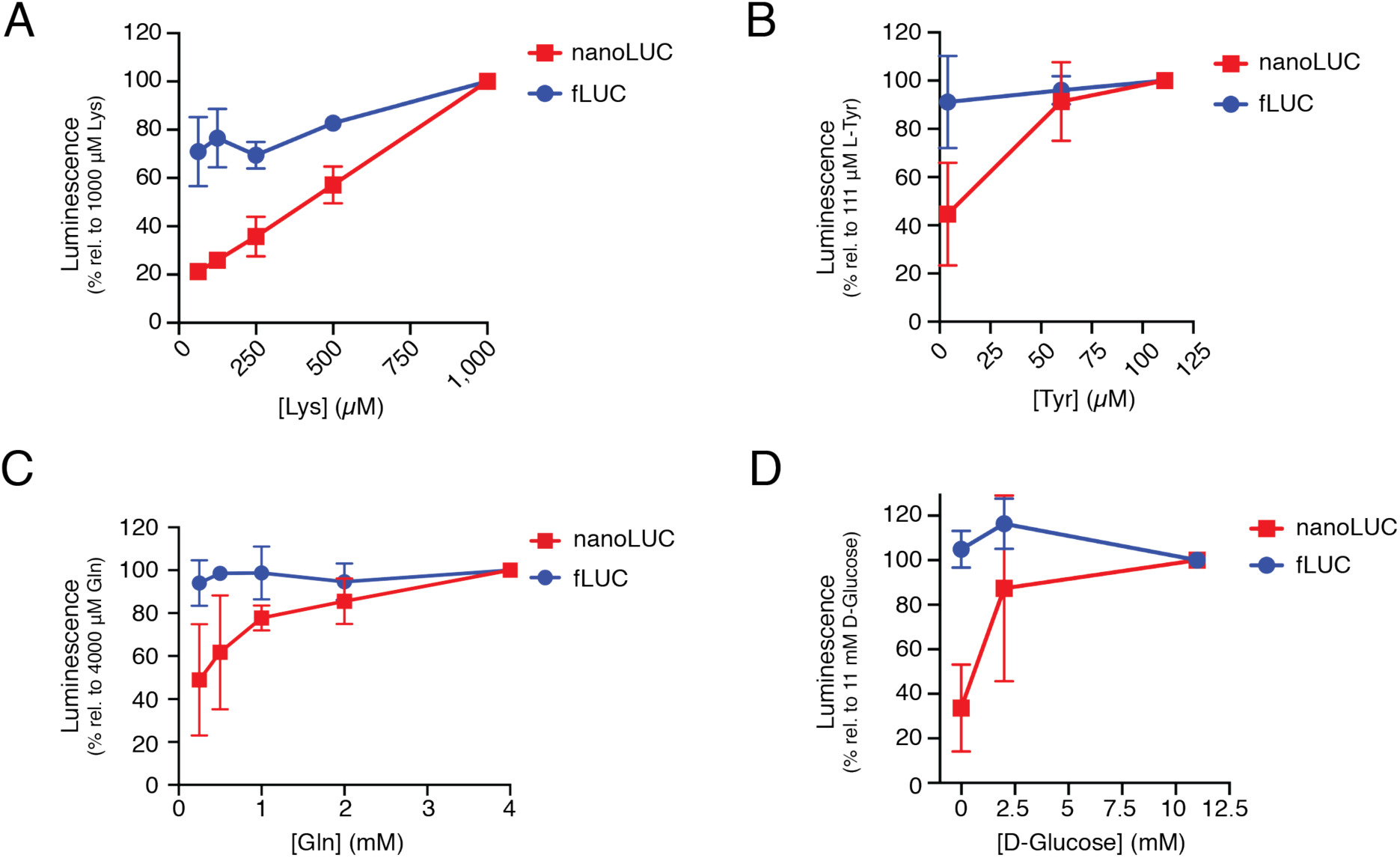
The 5’ region of *Tg*ApiAT1 mediates regulation in response to a range of nutrients. NanoLUC and fLUC luminescence readings in a parasite strain expressing nanoLUC from the *Tg*ApiAT1 5’ region (red) and fLUC from the *α*-tubulin (tub) 5’ region (blue), and grown at a range of (**A**) [Lys], (**B**) [Tyr], (**C**) [Gln], and (**D**) D-glucose. Luminescence is expressed as a percent of the luminescence at the highest tested concentration of each nutrient for both nanoLUC and fLUC measurements. Data points represent the mean ± SD of three independent experiments for each nutrient.

**Figure S4.**
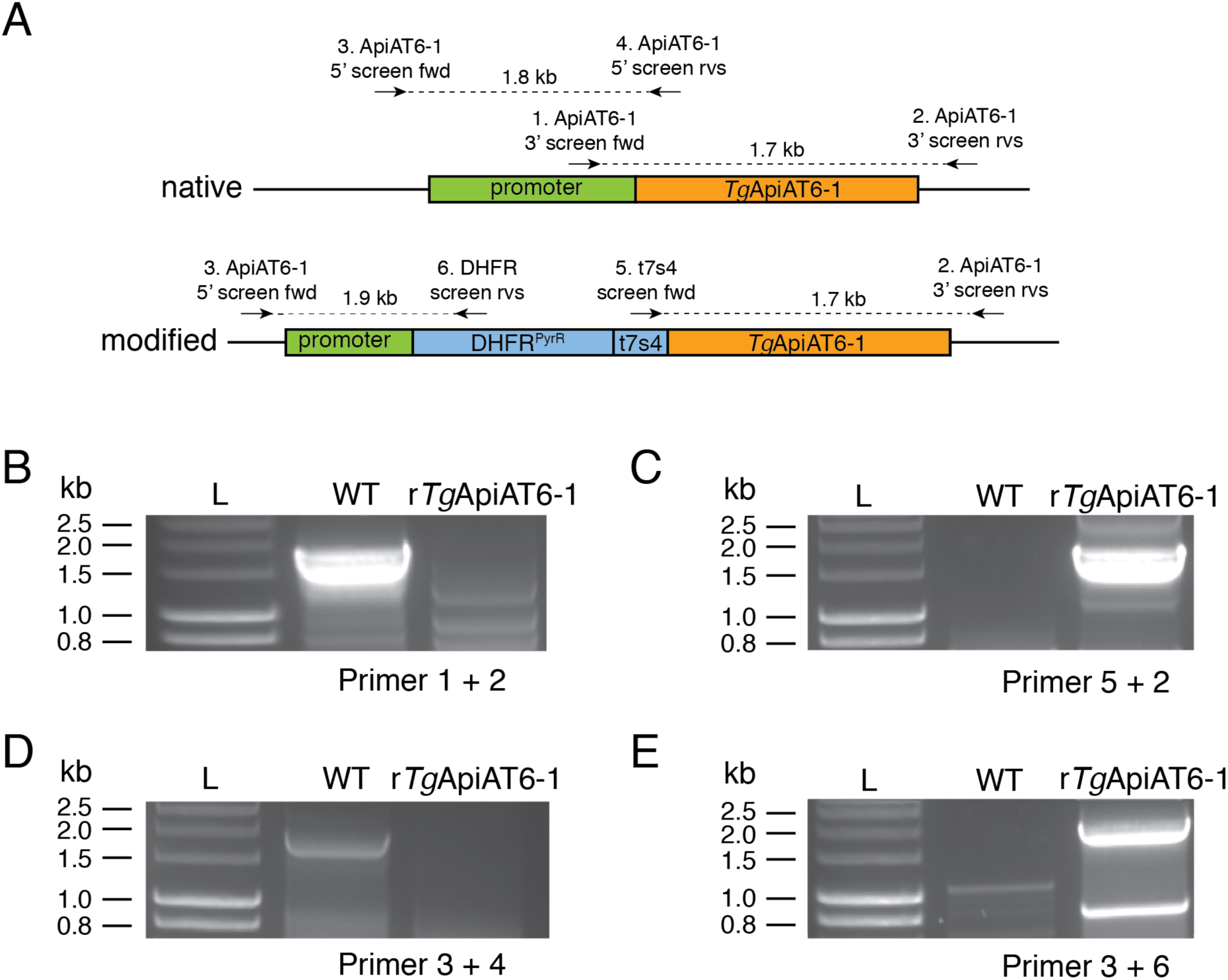
Generating an ATc-regulated *Tg*ApiAT6-1 strain. (**A**) Schematic depicting the promoter replacement strategy to generate the ATc-regulated *Tg*ApiAT6-1 strain (r*Tg*ApiAT6-1), and the positions of screening primers used in subsequent experiments to validate successful promoter replacement. The native locus (top) and promoter-replaced locus (bottom) are shown. DHFR^PyrR^, pyrimethamine-resistant dihydrofolate reductase cassette; t7s4, ATc-regulatable teto7-sag4 promoter. (**B-E**), PCR analysis using genomic DNA extracted from native RH strain (WT) and modified r*Tg*ApiAT6-1 strain parasites, with primers that specifically detect the 3’ region of the native locus (**B**), the 3’ region of the modified locus (**C**), the 5’ region of the native locus (**D**), and the 5’ region of the modified locus (**E**).

**Figure S5.**
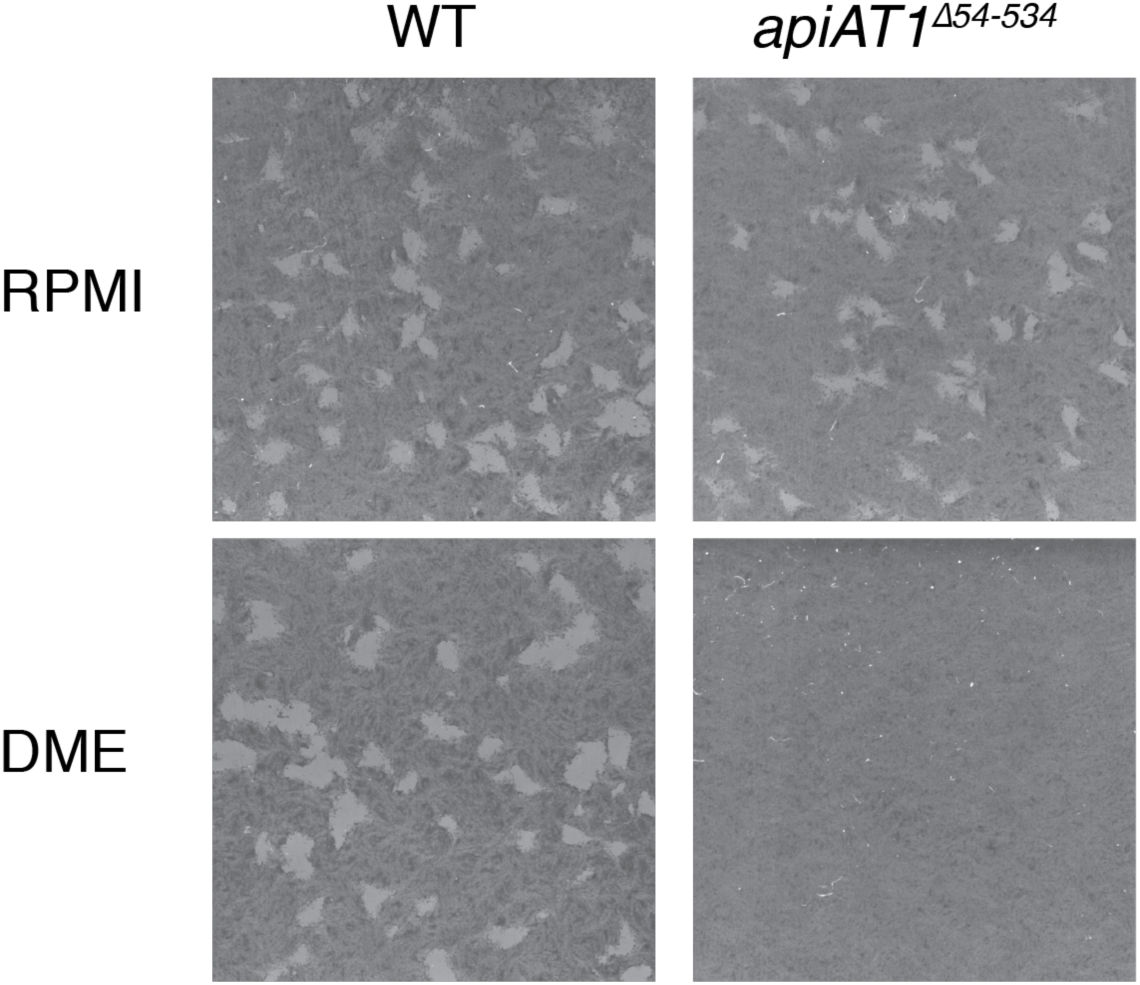
Disruption of *Tg*ApiAT1 impairs parasite growth in DME but not RPMI. 500 WT (RHΔ*hxgprt*/*apiAT1* 5’-nanoLUC/*tub*-fLUC; left) or *apiAT1*^Δ*54-534*^ (RHΔ*hxgprt*/*apiAT1* 5’-nanoLUC/*tub*-fLUC/*apiAT1*^Δ*54-534*^; right) parasites were inoculated into 25 cm^2^ tissue culture flasks containing either RPMI (top) or DME (bottom) and cultured for 8 days before staining with crystal violet to reveal plaque formation. Images are from a single experiment, and are representative of three independent experiments.

**Table S1. Data from the SWATH-MS proteomic analysis. Tab 1.** Averaged data from all replicates, indicating the ToxoDB ID, the number of peptides used in the analysis of each protein, *P* value, -log_10_ *P* value, the average fold change in the high vs low [Arg] conditions, the average fold change in the low vs high [Arg] conditions, the log_2_ fold change in the low vs high [Arg] condition, and the protein annotation. **Tab 2.** The data from each replicate of the experiment. H = 1.15 mM Arg; L = 50 µM Arg.

**Table S2.**
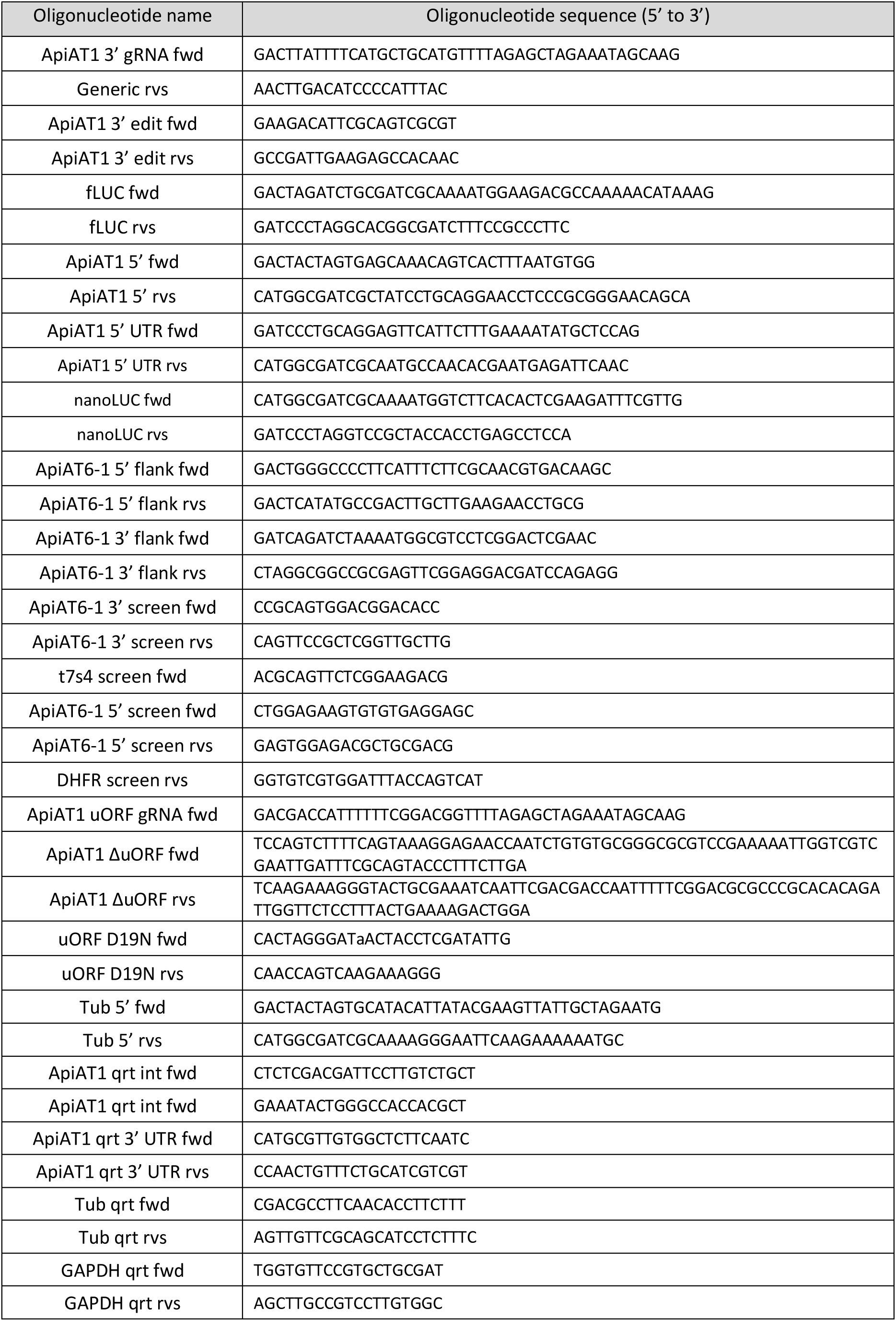
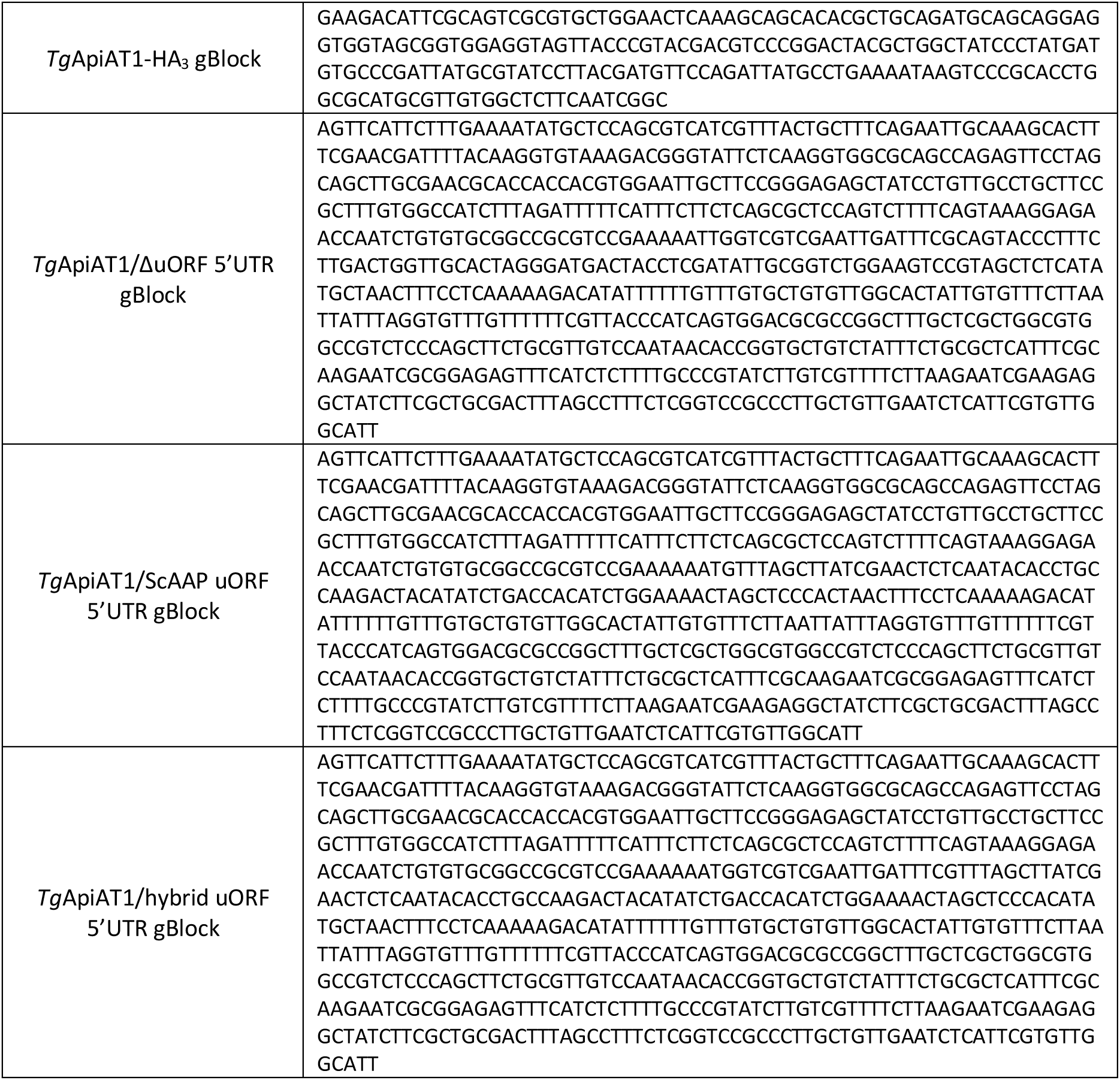
Sequences of the primers and gBlocks used in this study.

